# ClinOracle: Hierarchical AI Prediction of Target Binding and Patient-Derived Functional Activity Across Diverse Therapeutic Targets

**DOI:** 10.64898/2026.08.02.742378

**Authors:** Somaya A. Abdel-Rahman, Moustafa T. Gabr

## Abstract

AI platforms for drug discovery routinely achieve high hit rates against biochemical targets, yet the central translational challenge remains predicting whether a compound will be functionally active in patient-derived human cells. Here, we present ClinOracle, a hierarchical graph neural network that jointly predicts target binding and patient-derived functional activity by modeling functional activity as conditional on target engagement. Applied across five therapeutic targets spanning oncology, autoimmune, and neuroinflammatory diseases, ClinOracle ranked candidates using a Priority Score integrating translational probability with a developability score based on ADME and drug-likeness, advancing prioritized compounds through multistage prospective validation from biophysical binding to in vivo efficacy. Compounds with the highest Priority Scores consistently outperformed lower-ranked candidates across prospective experimental validation, demonstrating that hierarchical AI can prioritize compounds with patient-derived functional activity directly from molecular structure, rather than biochemical activity alone.

## INTRODUCTION

The central promise of AI-guided drug discovery, that computational prediction can meaningfully compress the timeline from target to therapy, remains largely unrealized at the translational level. Most AI-identified compounds are validated only through biochemical binding assays against purified proteins, which cannot determine whether a compound engages its target in living cells, elicits the intended functional response in disease-relevant biology, or remains active in patient-derived cells. This translational gap is reflected in the persistently high attrition rate of clinical drug development, where approximately 90% of candidates fail and nearly half of failures are attributed to insufficient efficacy rather than pharmacokinetic (PK) limitations (1–4). Our previous HTS-Oracle platform, like other AI platforms trained solely on biochemical binding, implicitly inherited this limitation (5,6). A more consequential challenge is to predict, directly from molecular structure, not only target binding but also functional activity in patient-derived cells. To our knowledge, no AI drug discovery platform has prospectively demonstrated this capability across multiple therapeutic targets. Here, we develop and prospectively validate such a framework.

Target engagement in intact cells is fundamentally different from, and more demanding than, measuring binding affinity. A compound may bind a purified extracellular domain with nanomolar affinity yet fail to access intracellular targets, be effluxed before reaching sufficient concentration, or engage a non-functional conformation absent in the native cellular context. The cellular thermal shift assay (CETSA) provides label-free confirmation of target engagement in intact cells and has been validated across kinases, epigenetic regulators, and immune targets (7–15). However, experimental confirmation alone cannot support prospective screening at the scale of hundreds of thousands of compounds.

A computational framework is therefore needed to predict, before experimental testing, which compounds are most likely to advance through successive stages of validation, including functional evaluation in patient-derived cells. Graph neural networks have emerged as powerful molecular representation frameworks for structure-activity learning, and multimodal architectures have further improved prospective hit rates across challenging target classes, including immune receptors and protein-protein interaction surfaces historically resistant to small molecule discovery (16–26). To bridge this translational gap, we developed ClinOracle, a multi-stage AI platform whose computational core employs a hierarchical multitask graph neural network that jointly predicts target binding probability and patient-derived functional activity. The ClinOracle GNN architecture comprises a shared molecular encoder and two prediction heads trained in a biologically grounded hierarchy: the first head estimates P(Binder) from molecular structure, while the second head estimates P(PBMC active | Binder) exclusively among experimentally confirmed binders, reflecting the biological reality that functional activity is downstream of and contingent upon target engagement. The integrated translational probability, P(Translationally active) = P(Binder) × P(PBMC active | Binder), provides a single compound-level score that optimizes for translational success rather than binding affinity alone.

The five targets selected for ClinOracle were chosen to test platform generalizability across disease areas unified by dysregulated immune signaling and the limited availability of small molecule modulators. The selected targets intentionally span extracellular immune checkpoints, intracellular kinases, innate immune signaling, and adaptive immune regulation to evaluate whether the hierarchical framework generalizes across distinct biological mechanisms rather than closely related proteins. LAG-3, a co-inhibitory checkpoint validated clinically by the FDA approval of relatlimab plus nivolumab (Opdualag), has no small molecule modulator beyond early preclinical stages (27–29). Siglec-15, an emerging checkpoint validated by the Phase I trial of NC318, remains similarly undrugged at the small molecule level (30–32). STING, the central innate immune DNA sensor and a well-established driver of disease activity in systemic lupus erythematosus (42), has an antagonist space that remains almost entirely unexplored despite intense clinical interest (33–35). RIPK1, a mediator of necroptosis and neuroinflammatory signaling with multiple clinical-stage inhibitors in Phase I/II trials, still lacks scaffolds with confirmed cellular engagement in primary neuroinflammatory systems (36–38). IRAK4, implicated in TLR4-driven microglial activation in Alzheimer’s disease (AD), remains comparatively unexplored at the small molecule level with no approved agent (39–41). Together, these five targets span T cell exhaustion, PD-L1-independent tumor immunosuppression, innate immune DNA sensing, and TLR- and necroptosis-driven neuroinflammation, three mechanistically distinct disease contexts providing a stringent test of platform generalizability.

ClinOracle demonstrates for the first time that an AI model can predict, from molecular structure alone, both target binding and functional activity in patient-derived primary human immune cells, with cross-disease generalization confirmed by held-out test set validation across five targets spanning three mechanistically distinct disease areas. Compound advancement was guided by a Priority Score combining this translational probability with a computational developability score reflecting ADME, drug-likeness, and synthetic accessibility, and the platform further advances AI-predicted hits through a mandatory sequential validation cascade encompassing biophysical binding confirmation, cellular target engagement, ADME profiling, patient-derived PBMC functional validation, and in vivo efficacy confirmation, establishing a replicable translational framework for AI-guided drug discovery.

## RESULTS

### ClinOracle: Platform Architecture and Target Selection

To address the translational gap between biochemical target binding and functional activity in patient-derived cells, we developed ClinOracle, a six-stage translational validation platform centered on a hierarchical graph neural network that jointly predicts target binding and patient-derived functional activity. ClinOracle integrates these predictions into a single ClinOracle Score, defined as P(Translationally active) = P(Binder) × P(PBMC active | Binder), which prioritizes compounds predicted to both bind their target and produce functional activity in patient-derived clinical samples. Prioritized compounds progress through mandatory biophysical, cellular, pharmacokinetic, patient-derived functional, and in vivo validation stages before candidate nomination (Figure 1A).

**Figure 1.**
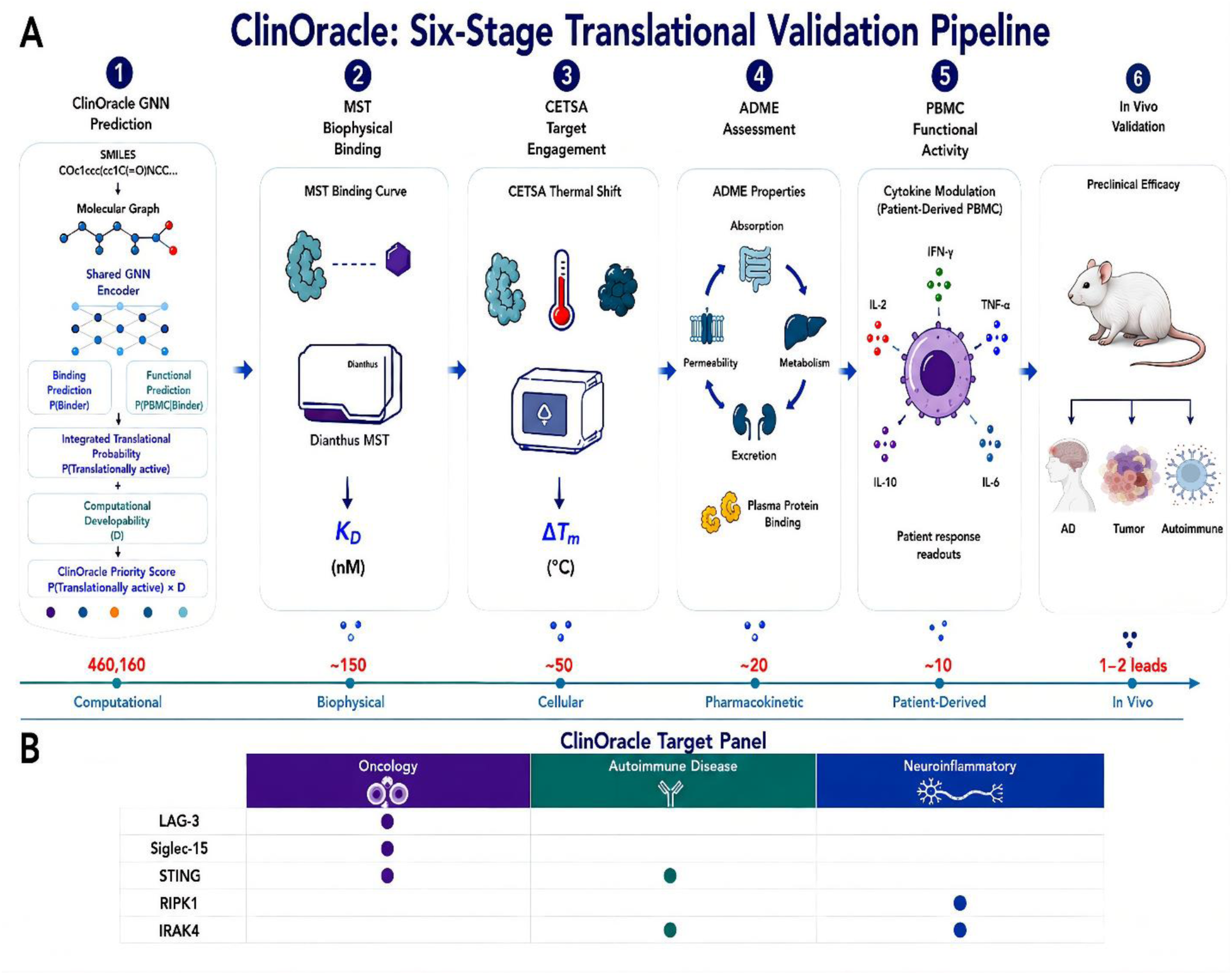
ClinOracle: An AI Platform for Predicting Functional Activity in Patient-Derived Clinical Samples Across Oncology, Autoimmune, and Neuroinflammatory Targets. **(A)** The ClinOracle six-stage translational validation pipeline implements mandatory sequential attrition from a 460,160-compound Enamine drug-like library through GNN-based virtual screening, in which candidates were ranked by a ClinOracle Priority Score combining the model’s translational probability, P(Translationally active), with a computational developability score reflecting drug-likeness and ADME (∼150 top-ranked hits per target), followed by biophysical binding confirmation by MST, cellular target engagement by CETSA, ADME profiling of top leads, patient-derived PBMC functional validation, and in vivo efficacy confirmation for the top lead(s) per target. **(B)** The ClinOracle target panel comprising five immunologically and therapeutically distinct targets, LAG-3, Siglec-15, STING, RIPK1, and IRAK4, spanning three mechanistically distinct disease areas: oncology, autoimmune, and neuroinflammatory disease.

Beginning with a GNN-based virtual screening engine applied prospectively to a library of 460,160 Enamine drug-like compounds, ClinOracle implements progressive attrition across six stages: ClinOracle-predicted hits (∼150 compounds, top 0.03% of library) are first subjected to biophysical binding confirmation by microscale thermophoresis (MST), reducing the hit set to compounds with experimentally confirmed target affinity. Confirmed binders are then advanced to cellular target engagement confirmation by CETSA, yielding compounds with demonstrated engagement in both model and patient-relevant cellular contexts. Top compounds are subsequently profiled for ADME properties including microsomal stability, Caco-2 permeability, and plasma protein binding. These leads are subjected to functional validation in disease-relevant patient-derived PBMC assays measuring target-specific immunological outputs. The most active compounds per target are advanced to in vivo efficacy testing in disease-relevant animal models. This mandatory sequential architecture (Figure 1A) ensures that no compound advances on the basis of computational prediction or binding affinity alone, directly addressing the primary mechanism by which AI-identified hits fail to translate to clinical efficacy.

ClinOracle was applied across the five-target, three-disease-area panel introduced above, oncology (LAG-3, Siglec-15), autoimmune disease (STING), and neuroinflammatory disease (RIPK1, IRAK4), selected to constitute a rigorous and deliberate test of platform generalizability for predicting binding and functional activity across diverse clinical contexts (Figure 1B).

Existing AI-enabled drug-discovery approaches address distinct components of the discovery process but generally do not model progression from molecular recognition to functional activity in patient-derived cells. Structure-based methods, including AlphaFold-guided screening and Boltz models, support structure prediction, molecular interaction modeling, and, in some implementations, affinity or developability prediction. AtomNet has demonstrated prospective hit identification across diverse targets, with selected predictions validated experimentally in biochemical assays. Our previous HTS-Oracle platform extended prospective AI-guided screening to immune-regulatory targets and achieved biophysically confirmed hit rates of ∼30%, representing substantial enrichment over unguided screening (5,6). However, these approaches primarily prioritize compounds on the basis of predicted molecular or target-level properties and do not explicitly model the conditional progression from target binding to functional activity in patient-derived cells. ClinOracle addresses this gap by integrating P(Binder) and P(PBMC active | Binder) into a unified translational-activity score. To our knowledge, ClinOracle is the first framework prospectively evaluated for this combined objective across multiple targets.

### ClinOracle’s hierarchical GNN jointly predicts target binding and patient-derived functional activity across five targets

To enable large-scale prospective identification of compounds with both target-binding and patient-derived functional activity, we developed a hierarchical multitask GNN comprising a shared molecular encoder and two sequentially dependent prediction heads (Figure 2A; see Methods). The two heads combine into P(Translationally active) = P(Binder) × P(PBMC active | Binder), the biological prediction generated by the GNN; compound nomination for testing additionally incorporates a computational developability score (D) to generate the ClinOracle Priority Score = P(Translationally active) × D. Models were trained independently for each of five targets using 25,000 experimentally screened compounds per target (Supplementary Table S1), with ΔF_norm_ ≥ 20% defining confirmed binders (QC pass rate >99.8%; Supplementary Table S2).

**Figure 2.**
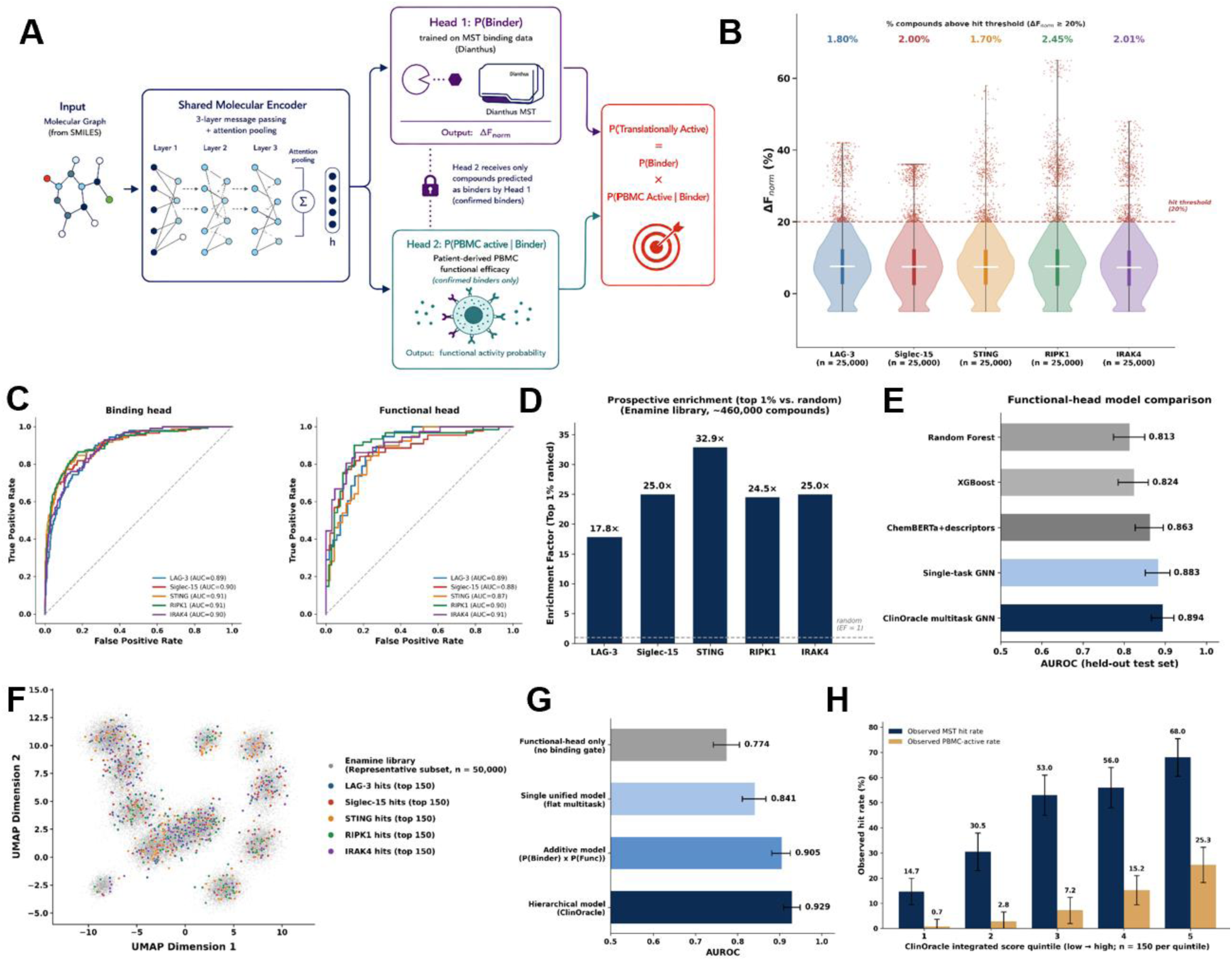
ClinOracle’s hierarchical multitask GNN jointly predicts target binding and patient-derived functional activity across five targets. **(A)** Architecture: a shared molecular encoder feeds two sequentially dependent heads, P(Binder) (trained on Dianthus MST binding data) and P(PBMC active | Binder) (trained exclusively on confirmed binders using patient-derived PBMC functional efficacy scores), combined into P(Translationally active) = P(Binder) × P(PBMC active | Binder). **(B)** Distribution of ΔF_norm_ values across the five MST screening datasets (n = 25,000 compounds per target); colored dots indicate confirmed hits (ΔF_norm_ ≥ 20%); hit rates ranged from 1.70% (STING) to 2.45% (RIPK1). **(C)** ROC curves for binding-head classification on a held-out test set (left; n = 5,000 compounds/target; AUC 0.89-0.91) and functional-head classification among confirmed binders (right; n = 85-127 compounds/target; AUC 0.87-0.91). **(D)** Prospective enrichment factors for the binding head at the top 1% cutoff, ranging from 17.8× (LAG-3) to 32.9× (STING). **(E)** Functional-head model comparison (AUROC, 95% CI) against Random Forest, XGBoost, ChemBERTa, and a single-task GNN; corresponding binding-head comparison shown in Figure S3. **(F)** UMAP visualization of chemical space; a representative subset of 50,000 compounds from the 460,160-compound Enamine Hit Locator Library (gray) alongside the top 150 ClinOracle-predicted hits per target (colored; 750 total), jointly embedded using Morgan fingerprints (radius = 2, 1,024 bits) and Jaccard distance. **(G)** Ablation comparing the hierarchical multiplicative integrated score against three alternatives on a shared held-out set of 3,500 compounds (background positive rate 5.7%): functional-head-only, single unified model, additive combination, and the hierarchical ClinOracle product; error bars show bootstrapped 95% confidence intervals (2,000 resamples), non-overlapping between adjacent variants except additive-versus-hierarchical (0.909-0.925 overlap). **(H)** Compounds selected using the ClinOracle Priority Score were stratified by P(Translationally active) into quintiles across the 750 compounds (150/target) advanced into the prospective screening cascade, showing observed MST binding hit rate and PBMC functional activity rate per quintile (n = 150/quintile; error bars, 95% Wilson confidence intervals).

Binding-head screening distributions were consistent across targets, with confirmed-binder rates from 1.70% (STING, 425/25,000) to 2.45% (RIPK1, 613/25,000) and median ΔF_norm_ of 7.28-7.56% (Figure 2B). Binding-head performance on a held-out test set (5,000 compounds/target) was strong and consistent, with ROC-AUC from 0.89 (LAG-3) to 0.91 (STING) (Figure 2C, left), and prospective enrichment at the top 1% cutoff from 17.8× (LAG-3) to 32.9× (STING) (Figure 2D). Precision-recall analysis, more diagnostic than ROC-AUC alone at this low background positive rate, confirmed the binding head substantially outperforms random prioritization (average precision 0.20-0.41 across targets; Figure S1).

Restricting evaluation to confirmed binders, the functional head predicted PBMC functional activity with AUROC from 0.87 (STING) to 0.91 (IRAK4), and predicted-versus-observed functional scores correlating at Pearson r = 0.605-0.654 across all five targets (all p < 0.0001; Figure 2C, right; Figure S2), yielding average precision of 0.84-0.89 across targets (baseline prevalence 44.4%; Figure S1). The functional head outperformed conventional non-graph baselines (Random Forest, mean AUROC 0.813; XGBoost, 0.824) and representation-learning/graph-based baselines (ChemBERTa-plus-descriptors, 0.863; single-task GNN, 0.883), versus 0.894 for the ClinOracle multitask GNN (Figure 2E; corresponding binding-head comparison in Figure S3).

The integrated P(Translationally active) score generalized robustly across targets, with per-target held-out test AUROC from 0.968 (Siglec-15) to 0.985 (LAG-3), AUPRC 0.438-0.615, and enrichment at the top 1% of 47.5×-61.1×. Pooling predictions across all five targets (background positive rate 0.87%) yielded AUROC = 0.976, AUPRC = 0.522, 78% precision among the top 50 pooled-ranked compounds, and 54.6-fold enrichment at the top 1%. Because the integrated score combines two independently trained heads, raw values are optimized for ranking rather than calibrated probability; Platt scaling substantially improved calibration (Brier score decreased from 0.0139 to 0.0058; expected calibration error decreased from 0.064 to 0.002) with discrimination fully preserved (Figure S4). The uncalibrated score was used for prioritization/enrichment throughout; the calibrated score provides an interpretable probability where needed. UMAP visualization confirmed ClinOracle-predicted hits (750 total, 150/target) span diverse chemical space rather than clustering (Figure 2F).

We next tested whether the hierarchical, multiplicative structure of the integrated score, rather than the presence of a GNN per se, accounts for its performance, comparing P(Translationally active) against three alternatives on a shared held-out set of 3,500 compounds (background positive rate 5.7%): a functional-head-only model, a single unified model, and an additive combination of the two heads (Figure 2G). Performance rose monotonically with structural fidelity to the underlying biology: functional head alone, AUROC = 0.774 (95% CI 0.742-0.804); unified model, 0.841 (0.811-0.867); additive combination, 0.905 (0.881-0.925); hierarchical product, 0.929 (0.909-0.948), with bootstrapped 95% CIs non-overlapping for all adjacent pairs except additive-versus-hierarchical, which overlapped (0.909-0.925). This supports the biological rationale that explicitly encoding functional activity as conditional on target engagement, rather than learning or approximating this dependency implicitly, yields the strongest predictor of translational success.

Finally, we tested whether the integrated score predicts outcomes in the actual prospective screening campaign. Compounds selected using the ClinOracle Priority Score were stratified by P(Translationally active) into quintiles across the 750 GNN-selected compounds (150/target; n = 150/quintile): observed MST binding hit rate rose monotonically from 14.7% in the lowest quintile to 68.0% in the highest, and observed PBMC functional activity rose from 0.7% to 25.3% (Figure 2H). Of the 750 compounds, 334 were confirmed MST binders, 138 CETSA-positive, and 77 PBMC-active, with top-quintile compounds disproportionately represented at every downstream stage. This demonstrates that P(Translationally active) remains strongly associated with downstream experimental success within the model-selected cohort, complementing the independent evaluation already provided by held-out test set metrics, and provides the computational foundation for the experimental validation cascade that follows.

### Biophysical validation of ClinOracle-predicted hits by MST and SPR

Prospective experimental validation of the top 150 ClinOracle-predicted candidates per target against the Enamine Hit Locator Library yielded 334 confirmed binders out of 750 compounds tested across all five targets (overall hit rate 44.5%) (Fig. 3A-E). Hit rates varied by target: RIPK1 achieved the highest hit rate (78/150, 52.0%), followed by STING (73/150, 48.7%), LAG-3 (64/150, 42.7%), IRAK4 (63/150, 42.0%), and Siglec-15 (56/150, 37.3%). The consistently high hit rates across all five mechanistically distinct targets confirm that the ClinOracle GNN models generalize effectively beyond their training distributions to prospectively identify novel binders from a large, structurally diverse compound library.

**Figure 3.**
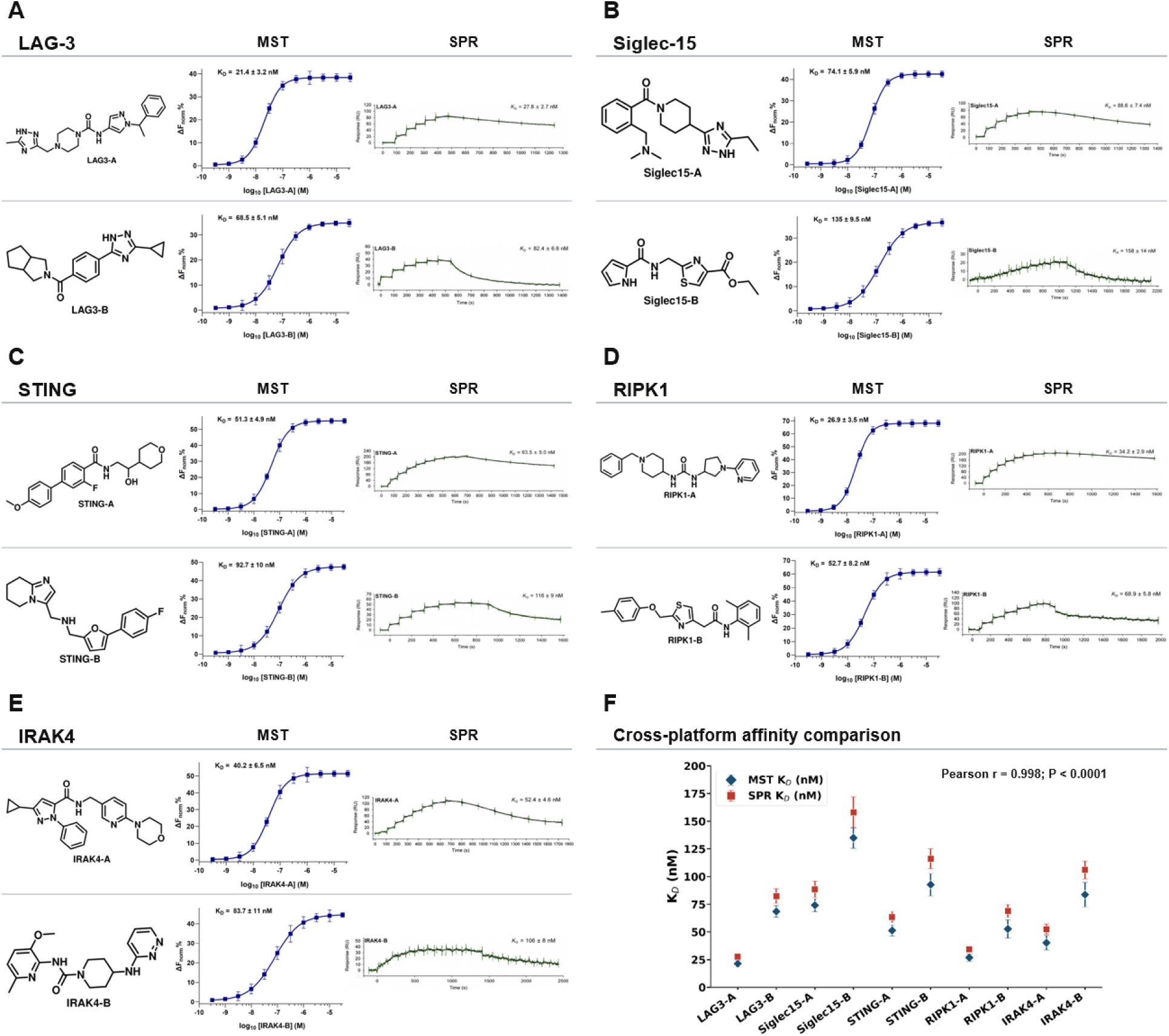
Biophysical validation of top two ClinOracle-predicted hits by MST and SPR across five targets. **(A-E)** Chemical structures, Dianthus MST binding curves, and SPR sensorgrams for the two top-ranked ClinOracle candidates per target: LAG-3 **(A)**, Siglec-15 **(B)**, STING **(C)**, RIPK1 **(D)**, and IRAK4 **(E)**. MST binding curves show ΔF_norm_ (%) as a function of ligand concentration (log scale); *K_D_* values were determined by sigmoidal dose-response fitting. SPR sensorgrams show binding response (RU) over time at multiple analyte concentrations; K_D_ values were determined by steady-state affinity analysis. **(F)** Binding affinity summary comparing MST (filled diamonds) and SPR (filled squares) K_D_ values for all 10 validated hits. Error bars represent mean ± SD (MST, n=5; SPR, n=3). MST and SPR K_D_ values were in close agreement across all compounds (within 1.5-fold), confirming orthogonal validation of ClinOracle-predicted binding affinities.

The two top-ranked ClinOracle candidates per target were advanced to orthogonal validation by surface plasmon resonance (SPR), confirmed MST-derived binding affinities. For LAG-3, compounds **LAG3-A** and **LAG3-B** demonstrated concentration-dependent sigmoidal binding curves by MST with *K_D_* values of 21.4 ± 3.2 nM and 68.5 ± 5.1 nM, respectively (Fig. 3A). SPR sensorgrams confirmed binding with *K_D_* values of 27.8 ± 2.7 nM (**LAG3-A**) and 82.4 ± 6.6 nM (**LAG3-B**), in close agreement with MST measurements. For Siglec-15, MST binding curves yielded *K_D_* values of 74.1 ± 5.9 nM (**Siglec15-A**) and 135 ± 9.5 nM (**Siglec15-B**), confirmed by SPR at 88.6 ± 7.4 nM and 158 ± 14 nM, respectively (Fig. 3B). For STING, hits **STING-A** and **STING-B** bound with MST *K_D_* values of 51.3 ± 4.9 nM and 92.7 ± 10.0 nM, confirmed by SPR at 63.5 ± 5.0 nM and 116 ± 9 nM, respectively (Fig. 3C). For RIPK1, hits **RIPK1-A** and **RIPK1-B** demonstrated MST *K_D_* values of 26.9 ± 3.5 nM and 52.7 ± 8.2 nM, confirmed by SPR at 34.2 ± 2.9 nM and 68.9 ± 5.8 nM, respectively (Fig. 3D). For IRAK4, hits **IRAK4-A** and **IRAK4-B** exhibited MST *K_D_* values of 40.2 ± 6.5 nM and 83.7 ± 11.0 nM, confirmed by SPR at 52.4 ± 4.6 nM and 106 ± 8.0 nM, respectively (Fig. 3E). Across all 10 validated compounds, MST and SPR *K_D_* values were in close agreement, with a maximum discordance of 1.5-fold, confirming the reliability of ClinOracle-predicted binding affinities (Fig. 3F). All 10 compounds exhibited *K_D_* values below 160 nM by both methods, with six compounds demonstrating sub-100 nM affinities. The most potent binders were identified for LAG-3 (**LAG3-A**, MST *K_D_* = 21.4 ± 3.2 nM) and RIPK1 (**RIPK1-A**, MST *K_D_* = 26.9 ± 3.5 nM).

### Cellular target engagement confirmed by CETSA

To confirm that ClinOracle-predicted hits engage their respective targets within the cellular environment, we employed dose-response CETSA, which measures the concentration-dependent stabilization of soluble target protein following compound treatment and heat challenge. All 10 validated hits were profiled across multiple concentrations, and sigmoidal dose-response curves were fitted to extract EC_50_ values as a measure of cellular target engagement potency.

All 10 compounds demonstrated clear concentration-dependent increases in normalized soluble target protein, confirming intracellular target engagement across all five targets (Figure 4). For LAG-3, **LAG3-A** showed the most potent cellular engagement (EC_50_ = 112 ± 7.3 nM), consistent with its superior binding affinity by MST and SPR, while **LAG3-B** demonstrated an EC_50_ of 331 ± 26 nM (Figure 4A-B). For Siglec-15, **Siglec15-A** and **Siglec15-B** exhibited EC_50_ values of 378 ± 32 nM and 692 ± 81 nM, respectively (Figure 4C-D). For STING, **STING-A** demonstrated an EC_50_ of 251 ± 33 nM and **STING-B** an EC_50_ of 498 ± 75 nM (Figure 4E-F). For RIPK1, **RIPK1-A** and **RIPK1-B** showed EC_50_ values of 145 ± 18 nM and 231 ± 25 nM, respectively (Figure 4G-H). For IRAK4, **IRAK4-A** demonstrated an EC_50_ of 193 ± 17 nM and **IRAK4-B** an EC_50_ of 501 ± 38 nM (Figure 4I-J). Across all 10 compounds, CETSA EC_50_ values ranged from 112 nM (**LAG3-A**) to 692 nM (**Siglec15-B**), confirming cellular target engagement at pharmacologically relevant concentrations. The rank order of cellular potency was broadly consistent with the rank order of binding affinities determined by MST and SPR, with the A-series compounds (**LAG3-A**, **RIPK1-A**, **IRAK4-A**, **STING-A**, **Siglec15-A**) consistently demonstrating superior cellular engagement compared to their B-series counterparts, in agreement with their lower K_D_ values.

**Figure 4.**
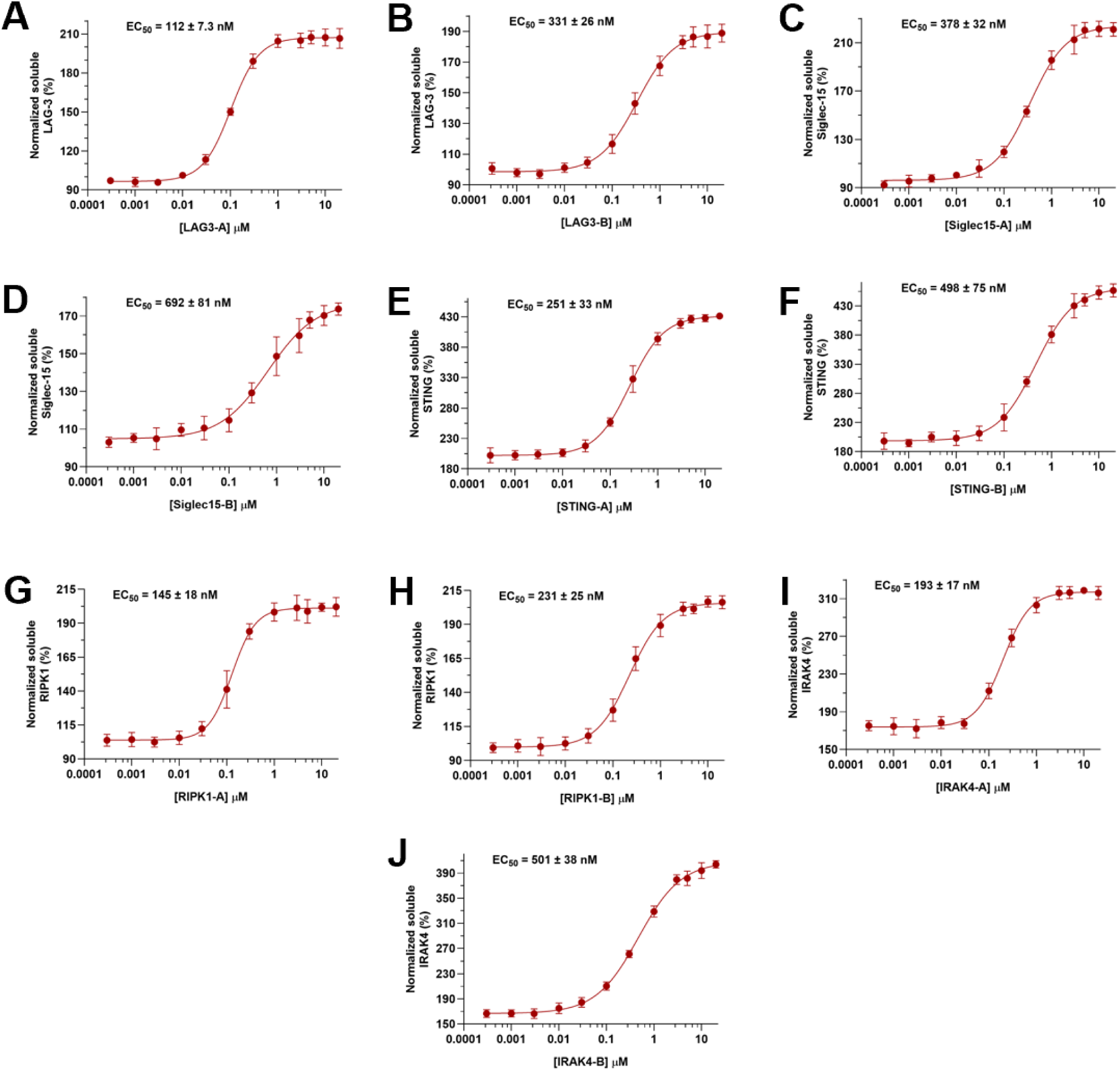
Cellular target engagement of ClinOracle-predicted hits by dose-response CETSA. **(A-B)** CETSA dose-response curves for LAG-3 hits **LAG3-A** and **LAG3-B**. **(C-D)** CETSA curves for Siglec-15 hits **Siglec15-A** and **Siglec15-B**. **(E-F)** CETSA curves for STING hits **STING-A** and **STING-B**. **(G-H)** CETSA curves for RIPK1 hits **RIPK1-A** and **RIPK1-B**. **(I-J)** CETSA curves for IRAK4 hits **IRAK4-A** and **IRAK4-B**. Y-axes show normalized soluble target protein (%) as a function of compound concentration (μM, log scale). Curves represent four-parameter sigmoidal dose-response fits; data points represent mean ± SD (n=5).

### ADME profiling confirms drug-like properties of ClinOracle-selected hits

A key objective of the ClinOracle pipeline is to identify compounds that not only bind their targets with high affinity but also possess favorable physicochemical and ADME properties suitable for downstream biological evaluation. This experimental characterization is distinct from the computational developability score used earlier to prioritize compounds for biophysical validation (see Materials and Methods); here, ADME properties were measured directly on the 10 validated hits following biophysical and cellular confirmation. To assess whether GNN-guided compound selection inherently enriches for drug-like hits, all 10 ClinOracle-validated compounds were profiled across a comprehensive in vitro ADME panel encompassing 19 parameters (Table 1).

**Table 1.**
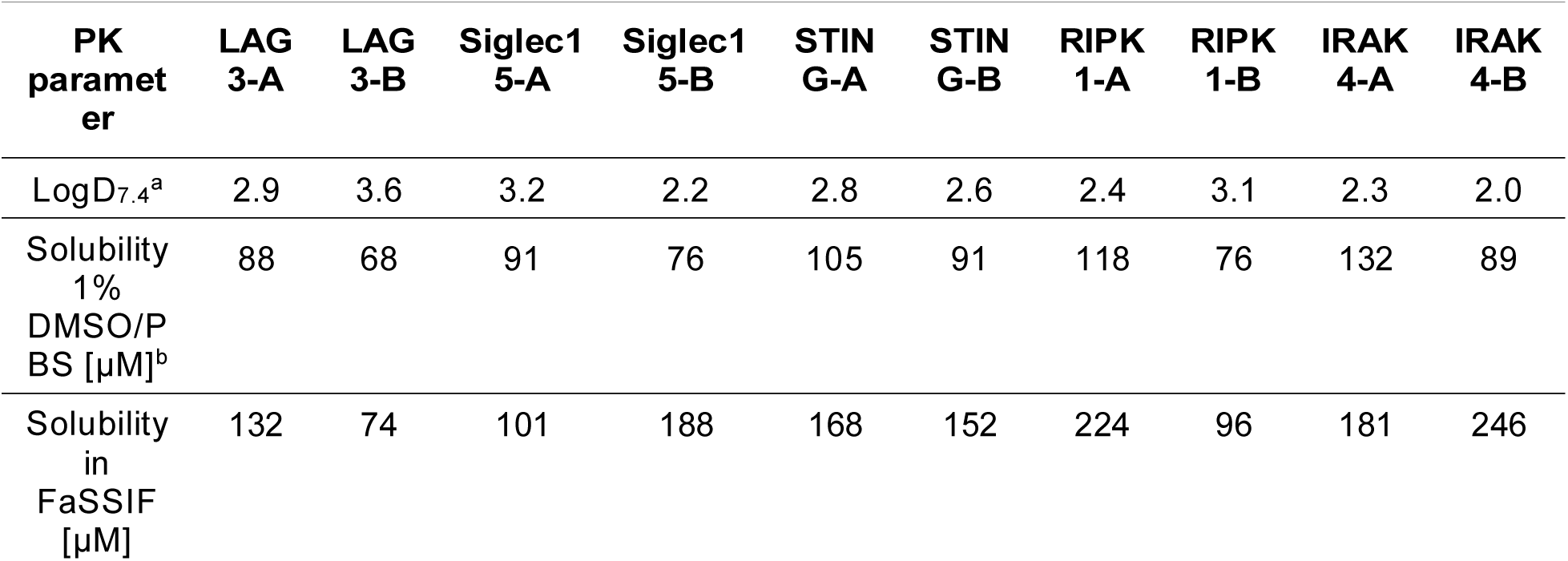
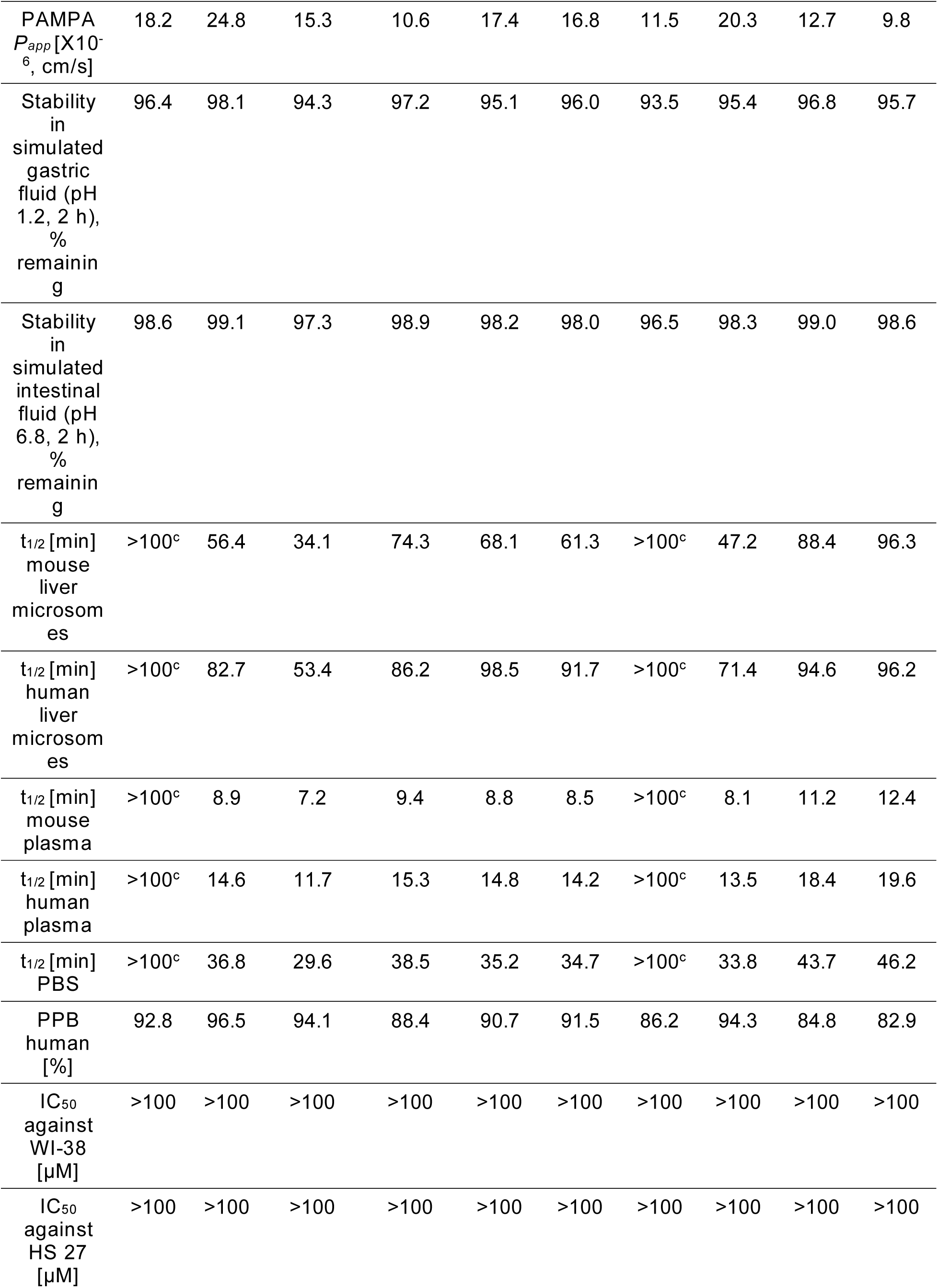
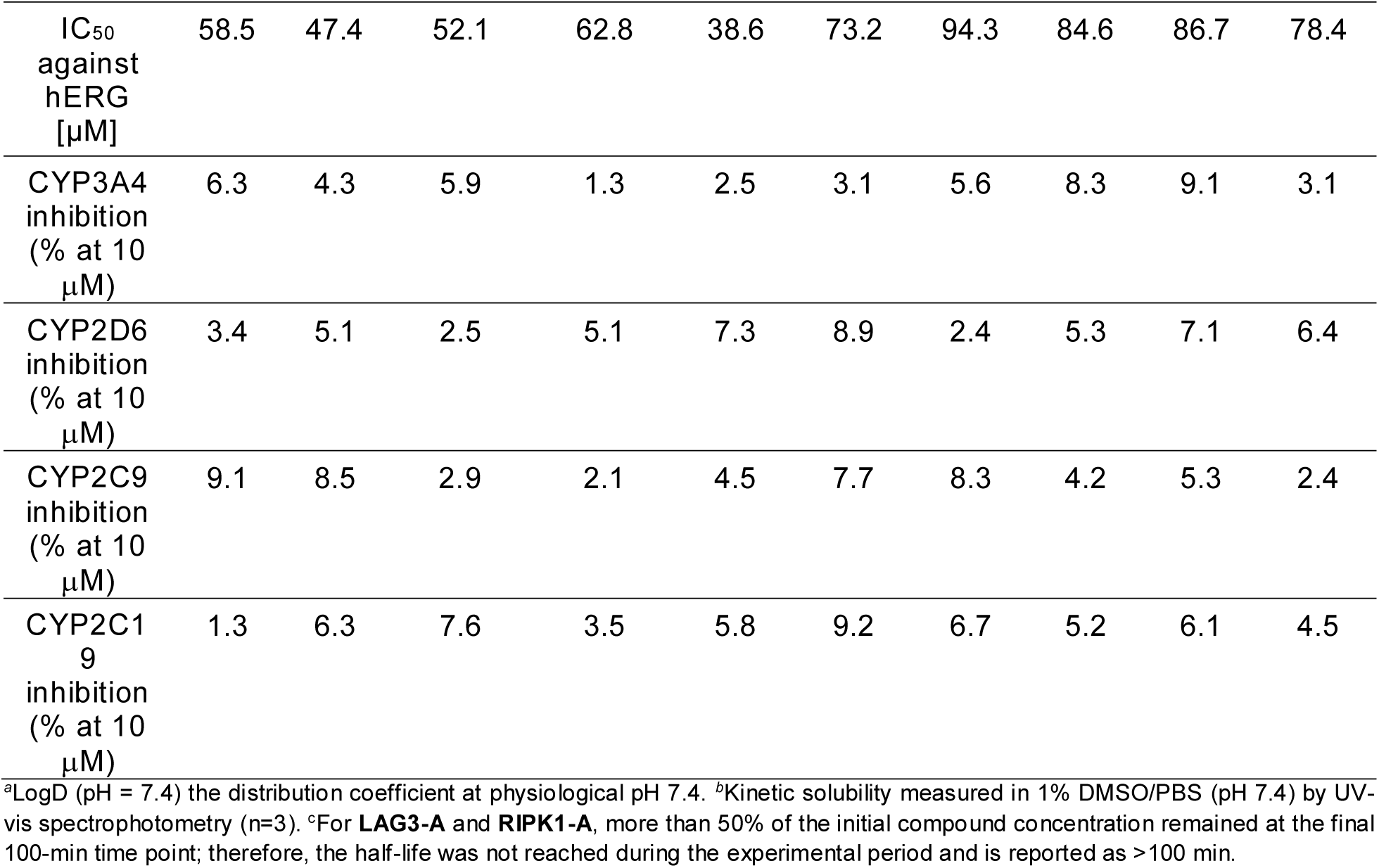
In vitro absorption, distribution, metabolism, and excretion (ADME) properties of the top 10 ClinOracle-derived hit compounds. Physicochemical properties, permeability, gastrointestinal stability, metabolic stability, plasma stability, plasma protein binding, cytotoxicity, hERG liability, and cytochrome P 450 inhibition profiles were evaluated to prioritize compounds for further experimental validation.

All 10 compounds exhibited LogD_7.4_ values of 2.0-3.6, kinetic aqueous solubility of 68-132 μM in 1% DMSO/PBS and 74-246 μM in FaSSIF, and passive PAMPA permeability of 9.8-24.8 × 10^−6^ cm/s, all within acceptable ranges for hit-stage compounds. Gastrointestinal stability was excellent across all 10 compounds, confirming chemical stability under physiologically relevant GI conditions. Excluding the two compounds whose half-lives exceeded the experimental observation period, human liver microsomal half-lives ranged from 53.4 to 98.5 min, whereas mouse liver microsomal half-lives ranged from 34.1 to 96.3 min. Aqueous stability varied across the series. **LAG3-A** and **RIPK1-A** exhibited PBS half-lives reported as >100 min, whereas the remaining compounds showed apparent half-lives of 29.6-46.2 min. Developability scores (D) are also presented for all 10 hits (D = 0.73-0.96; Supplementary Table S3), consistent with the experimental ADME profiles above.

Among the A-series lead compounds, **LAG3-A** and **RIPK1-A** demonstrated the most favorable metabolic stability profiles, with mouse and human liver microsomal half-lives reported as >100 min. **IRAK4-A** also showed favorable metabolic stability, with mouse and human microsomal half-lives of 88.4 and 94.6 min, respectively. **STING-A** and **Siglec15-A** exhibited human microsomal half-lives of 98.5 and 53.4 min, respectively. Plasma protein binding was high across all 10 compounds (82.9-96.5%), consistent with the lipophilic character of the hit series. **LAG3-A** and **RIPK1-A** exhibited the most favorable ex vivo stability, with apparent half-lives reported as >100 min in both mouse and human plasma. The remaining compounds showed shorter half-lives of 7.2-12.4 min in mouse plasma and 11.7-19.6 min in human plasma, identifying plasma stability as a potential lead-optimization liability for these compounds. The favorable plasma and microsomal stability of **LAG3-A** and **RIPK1-A** supported their advancement to in vivo PK evaluation.

Critically, all 10 ClinOracle hits demonstrated IC_50_ values exceeding 100 μM against both WI-38 and HS-27 normal human cell lines, confirming an absence of non-specific cytotoxicity at concentrations far exceeding their biologically active range and establishing a selectivity window of >1,000-fold relative to their MST K_D_ values. hERG inhibitory activity was minimal across all compounds (IC_50_ = 38.6-94.3 μM), providing a cardiac safety margin of >400-fold relative to binding affinities. CYP inhibition across four major isoforms (CYP3A4, CYP2D6, CYP2C9, CYP2C19) was negligible for all compounds at 10 μM (<10% inhibition in all cases), indicating a low risk of drug-drug interactions through CYP-mediated mechanisms.

### Patient-derived PBMC functional validation confirms disease-relevant activity of ClinOracle lead compounds

To determine whether ClinOracle-identified leads translate to functional activity in the disease contexts they are intended to address, the five A-series lead compounds were subjected to dose-response functional validation in patient-derived peripheral blood mononuclear cells (PBMCs) obtained from donors with confirmed diagnoses relevant to each target. PBMCs were sourced from BioIVT (n = 10 per disease cohort) and stimulated under target-specific conditions prior to compound treatment. Functional readouts were selected to capture the primary immunological consequence of each target’s modulation: T cell cytokine secretion for LAG-3 and Siglec-15, innate cytokine suppression for STING and IRAK4, necroptotic signaling suppression for RIPK1, and tumor cell killing for Siglec-15.

**LAG3-A** demonstrated concentration-dependent enhancement of T cell activation in colorectal cancer PBMCs, producing statistically significant increases in IFN-γ secretion (EC_50_ = 390 nM, E_max_ = 3.3-fold over vehicle) and IL-2 secretion (EC_50_ = 385 nM, E_max_ = 3.8-fold) across the full donor cohort (Figure 5A). Significance was first achieved at 30 nM for both readouts (p < 0.01), with maximal responses at 10,000 nM. The consistent dose-dependent activation across all 10 donors, despite inter-donor variability in baseline cytokine levels, confirms that **LAG3-A** restores T cell effector function in a patient-relevant colorectal cancer immune context.

**Figure 5.**
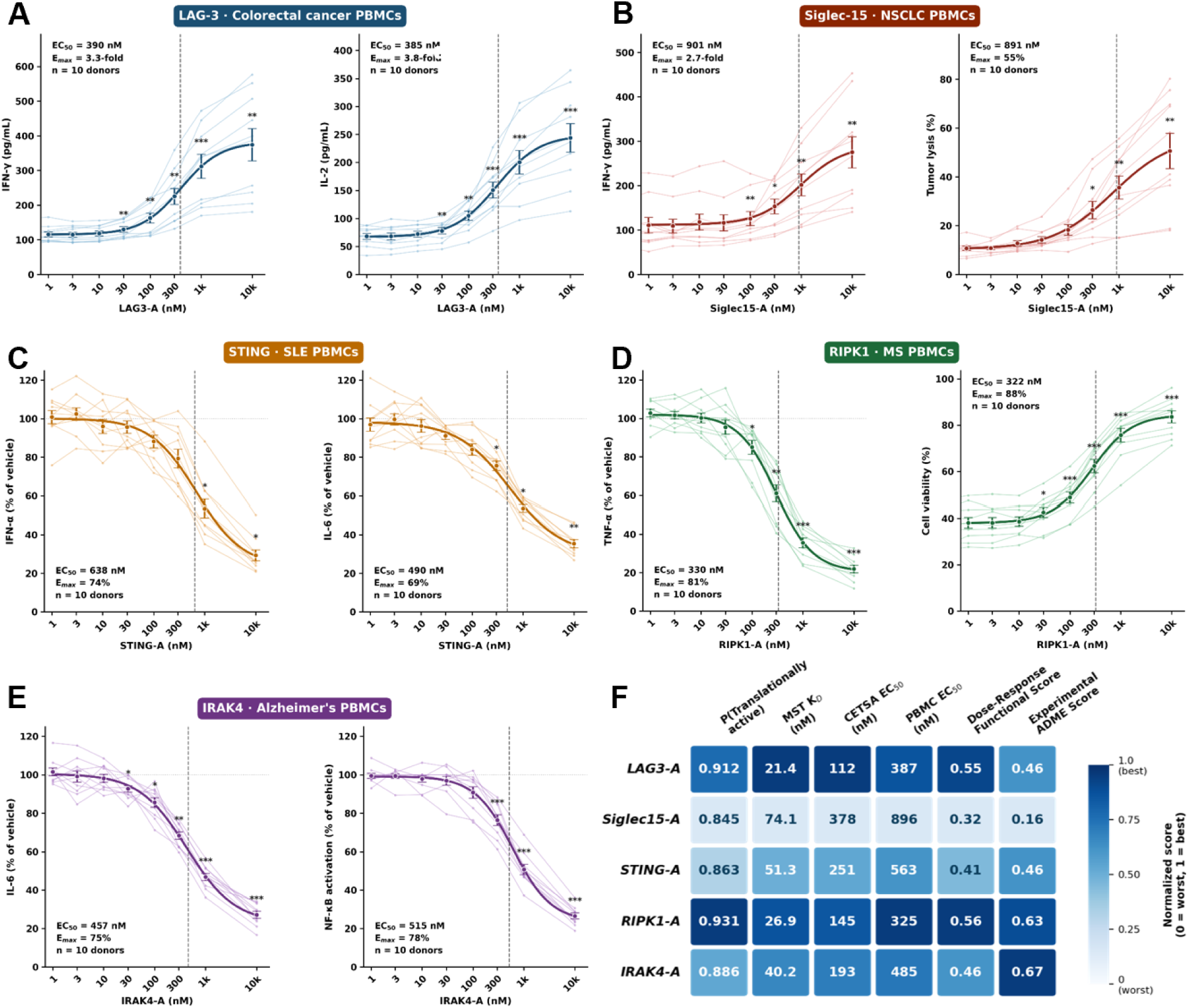
Patient-derived PBMC functional validation of ClinOracle-derived compounds across five immune and inflammatory targets. **(A)** Dose-response functional validation of **LAG3-A** in colorectal cancer patient-derived PBMCs (n = 10 donors). IFN-γ secretion (left) and IL-2 secretion (right) were measured across a concentration range of 1-10,000 nM. Individual donor traces represent per-donor means of three technical replicates. Bold curves show four-parameter logistic (4PL) fits to the mean data; error bars represent mean ± SEM across donors. Dashed vertical lines indicate EC_50_. Significance vs. vehicle: paired two-tailed t-test with Bonferroni correction across nine compound-versus-vehicle comparisons (k = 9). *p < 0.05, **p < 0.01, ***p < 0.001. **(B)** Dose-response functional validation of **Siglec15-A** in NSCLC patient-derived PBMCs (n = 10 donors). IFN-γ secretion (left) and tumor lysis (right, LDH release assay in co-culture with Siglec-15-expressing tumor cells at 10:1 effector-to-target ratio) were quantified across 1-10,000 nM. **(C)** Dose-response functional validation of **STING-A** in SLE patient-derived PBMCs (n = 10 donors). IFN-α suppression (left) and IL-6 suppression (right) are expressed as percentage of stimulated vehicle control per donor to account for inter-donor variability in baseline inflammatory tone. **(D)** Dose-response functional validation of **RIPK1-A** in MS patient-derived PBMCs (n = 10 donors). TNF-α suppression (left, normalized to stimulated vehicle) and cell viability recovery (right, absolute %) were measured across 1-10,000 nM. **(E)** Dose-response functional validation of **IRAK4-A** in AD patient-derived PBMCs (n = 10 donors). IL-6 suppression (left) and NF-κB activation suppression (right) are expressed as percentage of stimulated vehicle control per donor. **(F)** Integrated translational profile of the five ClinOracle lead compounds. Columns represent: P(Translationally active); MST K_D_; CETSA EC_50_; PBMC EC_50_; Dose-Response Functional Score (DFS; composite of normalized E_max_ and potency factor [1/(1 + log_10_(EC_50_/100))], range 0-1); and Experimental ADME Score (composite of human liver microsomal t½, PAMPA permeability, aqueous solubility, and hERG safety margin, normalized 0-1; Table 1). Color intensity reflects column-wise normalization (dark blue = best within column). P(Translationally active) is a model-predicted quantity; all other values are experimentally derived. Statistical analyses using paired two-tailed t-tests with Bonferroni correction for multiple comparisons.

**Siglec15-A** produced concentration-dependent enhancement of both T cell activation and tumor cytotoxicity in NSCLC patient PBMCs co-cultured with Siglec-15-expressing tumor cells (Figure 5B). IFN-γ secretion increased with an EC_50_ of 901 nM and E_max_ of 2.7-fold, while tumor lysis reached 55% at 10,000 nM (EC_50_ = 891 nM), representing a direct measure of restored antitumor cytotoxic activity. The parallel dose-dependence of cytokine secretion and tumor lysis supports that **Siglec15-A** functions through immune checkpoint modulation rather than direct cytotoxicity, consistent with its mechanism of action and the absence of cytotoxicity in normal cell lines (IC_50_> 100 μM; Table 1).

**STING-A** produced concentration-dependent suppression of pathological innate immune activation in systemic lupus erythematosus (SLE) patient PBMCs, with IFN-α suppression reaching 74% at 10,000 nM (EC_50_ = 638 nM) and IL-6 suppression reaching 69% (EC_50_ = 490 nM) (Figure 5C). Cytokine data are expressed as percentage of stimulated vehicle per donor to account for the substantial inter-donor variability in baseline inflammatory tone characteristic of SLE cohorts, which reflects the heterogeneity of disease activity at the time of donation. Statistically significant suppression was first observed at 1,000 nM for IFN-α and at 300 nM for IL-6, consistent with the compound’s CETSA EC_50_ of 251 nM and supporting on-target cellular engagement as the mechanism of observed suppression.

**RIPK1-A** demonstrated potent suppression of TNF-α secretion in multiple sclerosis (MS) patient PBMCs, achieving 81% suppression at 10,000 nM with an EC_50_ of 330 nM, and simultaneously restored cell viability in necroptosis-primed conditions to 88% of maximum at 10,000 nM (EC_50_ = 322 nM) (Figure 5D). The concordance between TNF-α suppression and viability rescue across the same concentration range, with significance emerging at 100 nM for TNF-α suppression and 30 nM for viability, confirms dual engagement of the RIPK1-mediated neuroinflammatory and necroptotic axes in a primary MS immune context. This dual functional profile distinguishes **RIPK1-A** as the lead with the highest translational relevance for neuroinflammatory disease.

**IRAK4-A** produced concentration-dependent suppression of TLR-driven innate immune activation in AD patient PBMCs, with IL-6 suppression reaching 75% at 10,000 nM (EC_50_ = 457 nM) and NF-κB activation suppression reaching 78% (EC_50_ = 515 nM) (Figure 5E). Significant suppression was observed from 30 nM for IL-6 and from 100 nM for NF-κB activation, consistent with the CETSA EC_50_ of 193 nM. The parallel suppression of both upstream cytokine production and downstream transcriptional activation consistent with on-target IRAK4 engagement as the mechanism of functional activity in AD-relevant immune cells.

Across all five targets, EC_50_ values in patient-derived PBMCs ranged from 325 to 896 nM, while CETSA EC_50_ values ranged from 112 to 378 nM, consistent with the additional barriers to target engagement in complex primary immune cell systems compared to cellular thermal shift conditions. The integrated translational profile (Figure 5F) demonstrates that all five lead compounds successfully traverse the full ClinOracle validation cascade from ClinOracle-predicted binding through biophysical confirmation, cellular target engagement, ADME profiling, and patient-derived functional activity.

### LAG3-A exhibits favorable PK properties, cross-species LAG-3 engagement, and antitumor efficacy in CT26 tumors

To determine whether **LAG3-A** achieved systemic exposure sufficient to support in vivo efficacy studies, PK analysis was performed in mice following single-dose intravenous and oral administration. Following intravenous administration at 2 mg/kg, **LAG3-A** exhibited a plasma half-life of 6.1 h, and systemic clearance of 3.9 mL/min/kg. The corresponding plasma exposure was AUC₀–∞ = 21.6 µM·h.

Following oral administration at 25 mg/kg, **LAG3-A** was efficiently absorbed, reaching a plasma C_max_ of 8.7 µM at a T_max_ of 1.5 h. Total systemic exposure following oral dosing reached AUC₀–∞ = 118.4 µM·h, corresponding to an estimated oral bioavailability of approximately 44%. The apparent plasma half-life following oral dosing was 6.8 h, supporting sustained systemic exposure over the dosing interval. Importantly, plasma concentrations achieved following oral administration remained above the biochemical binding potency and cellular functional activity range for a substantial portion of the 24 h dosing interval. These PK properties supported selection of 25 mg/kg once-daily oral dosing for subsequent efficacy testing in the CT26 syngeneic colon carcinoma model. Notably, **LAG3-A** was well tolerated at 25 mg/kg QD, with no significant changes in body weight observed throughout the treatment period.

Before evaluating antitumor activity, we confirmed that **LAG3-A** retained cross-species binding activity toward murine LAG-3. MST analysis demonstrated direct interaction between **LAG3-A** and recombinant mouse LAG-3, yielding a K_D_ of 30.2 ± 4.6 nM (Figure S5), comparable to the affinity observed for human LAG-3. This confirmed that **LAG3-A** engages murine LAG-3 with high affinity and supported evaluation in immunocompetent syngeneic mouse tumor models.

We next investigated the therapeutic activity of **LAG3-A** in the CT26 syngeneic colorectal carcinoma model in BALB/c mice. CT26 cells were implanted subcutaneously, and treatment was initiated once tumors reached approximately 75-100 mm^3^. Mice were randomized to receive vehicle, **LAG3-A** at 25 mg/kg by oral gavage once daily, anti-mouse PD-1 antibody, or the combination of **LAG3-A** and anti-PD-1 for 21 consecutive days.

Vehicle-treated tumor volume reached approximately 1,940 mm^3^ by Day 21 (Figure 6A). Anti-PD-1 monotherapy delayed tumor progression (∼37% growth inhibition relative to vehicle), **LAG3-A** monotherapy produced a stronger antitumor effect (∼58% growth inhibition), and combination treatment produced the greatest efficacy (∼75% growth inhibition). Thus, **LAG3-A** suppressed CT26 tumor progression as a single agent and further enhanced the therapeutic activity of PD-1 blockade.

**Figure 6.**
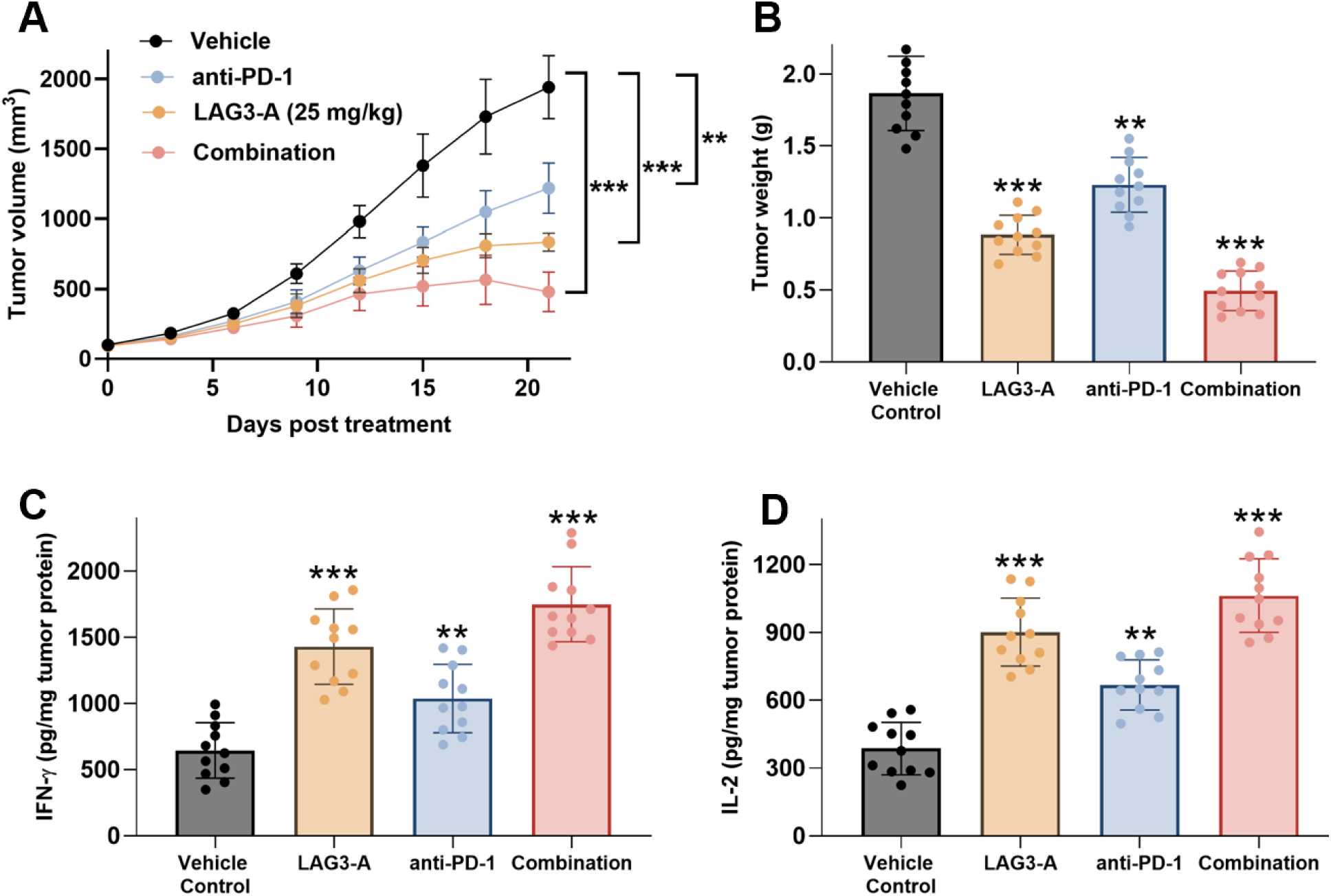
LAG3-A suppresses CT26 tumor growth and enhances the antitumor activity of PD-1 blockade in vivo. **LAG3-A** was selected for in vivo efficacy testing based on favorable mouse PK properties following oral dosing at 25 mg/kg. **(A)** CT26 tumor growth curves in BALB/c mice treated with vehicle, **LAG3-A** (25 mg/kg, PO, QD), anti-mouse PD-1 antibody, or the combination of **LAG3-A** and anti-PD-1. **(B)** Final tumor weights at study termination. **(C,D)** Intratumoral IFN-γ and IL-2 levels measured by ELISA from tumor lysates. Data are shown as mean ± SEM with individual animals shown where applicable. Statistical significance was assessed by two-way ANOVA for tumor growth curves and one-way ANOVA with multiple-comparison correction for endpoint analyses. **p < 0.01, ***p < 0.001.

Endpoint tumor weights closely paralleled the longitudinal tumor-volume measurements across all treatment groups (Figure 6B), confirming superior antitumor efficacy with combined LAG-3 and PD-1 blockade.

To determine whether tumor growth suppression was associated with enhanced antitumor immune activation, tumor lysates were analyzed by ELISA. **LAG3-A** markedly increased intratumoral IFN-γ and IL-2 levels relative to vehicle (Figure 6C,D); anti-PD-1 monotherapy produced smaller increases in both cytokines, while combination treatment produced the highest levels of each, consistent with enhanced antitumor immune activation.

### RIPK1-A demonstrates favorable PK properties, efficient brain penetration, and therapeutic efficacy in the 5xFAD model of AD

Before evaluating cognitive and neuroinflammatory efficacy, we confirmed that **RIPK1-A** retained cross-species binding activity toward murine RIPK1. MST analysis demonstrated direct interaction between **RIPK1-A** and recombinant mouse RIPK1, yielding a K_D_ of 15.4 nM (Figure S6), comparable to the affinity observed for human RIPK1 (MST K_D_ = 26.9 nM). This confirmed that **RIPK1-A** engages murine RIPK1 with high affinity and supported evaluation in the 5xFAD mouse model of AD. To determine whether **RIPK1-A** possessed PK properties compatible with in vivo efficacy studies, single-dose PK profiling was performed in male C57BL/6J mice (8-10 weeks old; n = 5) following intravenous and oral administration. After intravenous dosing (2 mg/kg), **RIPK1-A** exhibited a plasma elimination half-life of 5.4 h, a systemic clearance of 5.6 mL/min/kg, and a plasma AUC₀– ∞ of 15.6 µM·h. These parameters indicate low systemic clearance together with substantial tissue distribution, consistent with favorable in vivo drug disposition.

Following oral administration at 25 mg/kg, **RIPK1-A** was rapidly absorbed, reaching a plasma C_max_ of 9.2 µM at a T_max_ of 1.0 h. Total plasma exposure increased to an AUC₀– ∞ of 91.3 µM·h, corresponding to an estimated oral bioavailability of approximately 47%. The apparent terminal plasma half-life following oral dosing was 6.2 h, providing sustained systemic exposure throughout the dosing interval. Importantly, systemic exposure exceeded the biochemical binding affinity and cellular target-engagement concentrations throughout most of the dosing interval, supporting once-daily oral dosing. Because effective modulation of RIPK1 in AD requires adequate central nervous system exposure, brain PK exposure was also evaluated. **RIPK1-A** readily crossed the blood-brain barrier, achieving a brain C_max_ of 6.8 µM at 1 h and a brain AUC₀–∞ of 63.5 µM·h. The corresponding brain-to-plasma exposure ratio (K_p_) was 0.70, while correction for the measured unbound fractions yielded an estimated K_p,uu_ of 0.64, demonstrating efficient free-drug distribution into the brain.

We next evaluated the therapeutic efficacy of **RIPK1-A** in the 5xFAD transgenic mouse model of AD. Four-month-old male and female 5xFAD mice, together with age-matched wild-type littermates, were randomized into treatment groups (n = 12 per group; six males and six females) and treated with **RIPK1-A** (25 mg/kg, PO, QD) or vehicle for 21 consecutive days. This treatment window was selected because 5xFAD mice at this age exhibit established amyloid pathology, neuroinflammation, and measurable cognitive impairment while remaining responsive to pharmacological intervention.

Cognitive performance was first evaluated using the Y-maze spontaneous alternation test (Figure 7A). Vehicle-treated 5xFAD mice demonstrated significantly impaired spontaneous alternation compared with wild-type controls, reflecting deficits in hippocampal-dependent working memory. Administration of **RIPK1-A** significantly improved spontaneous alternation performance, restoring cognitive function toward wild-type levels. In contrast, no significant differences in total arm entries were observed among the experimental groups (Figure 7B), indicating that the improved cognitive performance was not attributable to alterations in locomotor activity or exploratory behavior.

**Figure 7.**
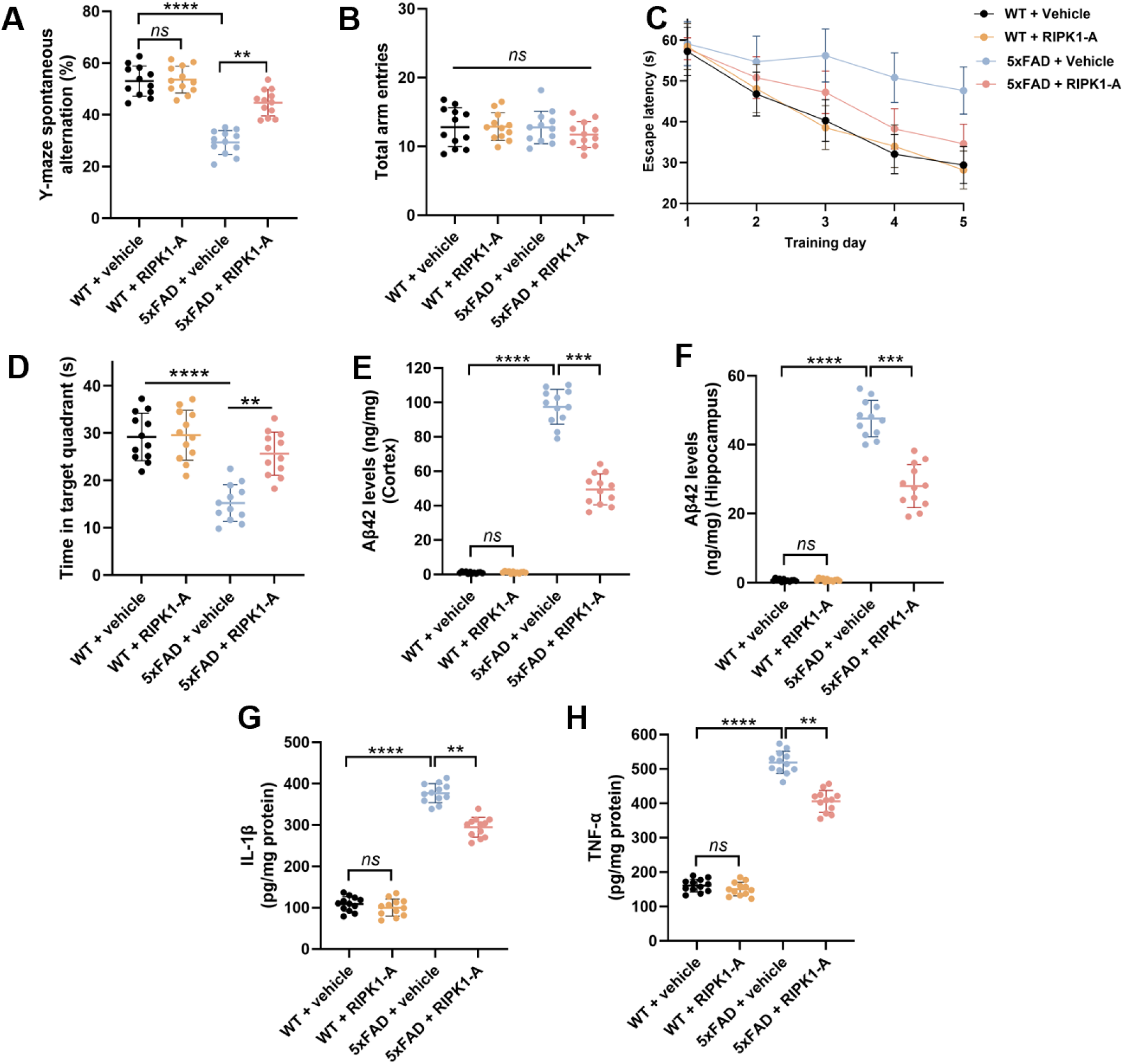
RIPK1-A improves cognitive function and attenuates AD pathology in 5xFAD mice. **(A)** Spontaneous alternation performance in the Y-maze demonstrating impaired working memory in vehicle-treated 5xFAD mice compared with wild-type (WT) littermates. Oral administration of **RIPK1-A** significantly restored spontaneous alternation toward WT levels. **(B)** Total arm entries recorded during the Y-maze test. Comparable exploratory activity among all experimental groups indicates that the cognitive improvements produced by **RIPK1-A** were not associated with alterations in locomotor behavior. **(C,D) RIPK1-A** improves spatial learning and memory in 5xFAD mice in the Morris water maze. **(C)** Escape latency during the acquisition phase. Vehicle-treated 5xFAD mice exhibited persistently longer escape latencies than WT controls, indicating impaired spatial learning. **RIPK1-A** treatment progressively improved learning performance in 5xFAD mice, resulting in shorter escape latencies during the final training sessions, whereas **RIPK1-A** had no appreciable effect in WT mice. **(D)** Spatial-memory retention during a 60-s probe trial, measured as time spent in the target quadrant. Vehicle-treated 5xFAD mice performed near chance level, whereas **RIPK1-A** treatment significantly increased target-quadrant occupancy. **(E,F)** Quantification of soluble Aβ42 in cortical **(E)** and hippocampal **(F)** tissue homogenates. **RIPK1-A** treatment significantly reduced amyloid burden in both brain regions relative to vehicle-treated 5xFAD mice. **(G,H)** Brain concentrations of the pro-inflammatory cytokines IL-1β **(G)** and TNF-α **(H)**. **RIPK1-A** markedly suppressed neuroinflammatory cytokine production in 5xFAD mice, consistent with inhibition of pathological inflammatory signaling. Data are presented as mean ± SD (n = 12 mice per group). Statistical analysis was performed using one-way ANOVA followed by Tukey’s multiple-comparison test. ns, not significant; *p < 0.05; **p < 0.01; ***p < 0.001; ****p < 0.0001.

The effects of **RIPK1-A** on spatial learning and memory were further assessed using the Morris water maze (Figure 7C,D). During the acquisition phase, vehicle-treated 5xFAD mice exhibited prolonged escape latencies compared with wild-type animals, consistent with impaired spatial learning. **RIPK1-A**-treated mice progressively improved throughout the training period, exhibiting significantly shorter escape latencies during the final training sessions. During the probe trial, vehicle-treated 5xFAD mice spent substantially less time in the target quadrant than wild-type controls, whereas **RIPK1-A** treatment significantly increased target quadrant occupancy, demonstrating improved spatial memory retention. Performance of wild-type animals was not significantly affected by **RIPK1-A** administration.

To determine whether the behavioral improvements were accompanied by changes in AD pathology, soluble Aβ42 concentrations were quantified in cortical and hippocampal tissue homogenates. Vehicle-treated 5xFAD mice exhibited marked accumulation of Aβ42 in both brain regions relative to wild-type littermates (Figure 7E,F). **RIPK1-A** treatment significantly reduced Aβ42 levels in both cortex and hippocampus, producing reductions of approximately 46% and 41%, respectively. Although these biochemical analyses demonstrate a substantial reduction in amyloid burden, future studies incorporating immunohistochemical plaque quantification will be important to determine whether RIPK1 inhibition also alters plaque deposition, morphology, and clearance kinetics.

Because RIPK1 signaling contributes directly to neuroinflammatory responses, inflammatory cytokine production was next examined. Vehicle-treated 5xFAD mice displayed significantly elevated concentrations of IL-1β and TNF-α, whereas treatment with **RIPK1-A** markedly suppressed both inflammatory mediators (Figure 7G,H), consistent with attenuation of pathological neuroinflammation in vivo.

To further characterize the effects of RIPK1 inhibition on microglial activation, flow cytometric analysis was performed using CD86 as a marker of pro-inflammatory microglia. Vehicle-treated 5xFAD mice exhibited a marked expansion of the CD86-positive microglial population relative to wild-type animals, whereas **RIPK1-A** significantly reduced this activated microglial subset (Figure S7).

## DISCUSSION

ClinOracle demonstrates that a hierarchical AI architecture explicitly modeling the dependency of functional activity on target engagement outperforms simpler formulations and translates into real experimental success across five structurally and biologically distinct targets. The ablation comparisons in Figure 2G show that this improvement is not incidental: performance increased monotonically across the four architectures, from a functional-head-only model (AUROC = 0.774) through unified and additive formulations to the full hierarchical architecture (AUROC = 0.929). Confidence intervals were non-overlapping for the earlier adjacent comparisons but overlapped between the additive and hierarchical models; therefore, the latter comparison supports a directional performance advantage rather than definitive separation based on confidence intervals alone. This progression indicates that encoding the biological dependency of function on binding explicitly, rather than approximating it implicitly, is itself responsible for the gain in predictive performance, not simply the use of a graph neural network. By validating these predictions through a cascade spanning biophysical, cellular, patient-derived, and in vivo endpoints, this work establishes a standard of translational evidence rarely achieved by computational drug discovery platforms, most of which terminate at binding confirmation without testing whether predicted hits translate into functional or therapeutic outcomes.

A key architectural feature of ClinOracle is the separation between biological prediction and compound prioritization. P(Translationally active) represents the trained, calibrated, and prospectively validated output of the hierarchical GNN, whereas the ClinOracle Priority Score additionally incorporates a computational developability score to guide practical compound selection. This distinction matters because binding and functional predictions and drug-likeness are not interchangeable properties; a compound can be highly likely to engage its target and function in patient cells while remaining a poor starting point for medicinal chemistry optimization, or vice versa. Collapsing these into a single score would obscure which failures reflect incorrect biological prediction and which reflect a deliberate developability trade-off, complicating both model evaluation and iterative improvement. This two-layer design mirrors how AI-guided discovery is organized in industrial settings, pairing a prediction engine that estimates biological success with a decision layer that balances biology against developability and other practical considerations. It also provides a natural framework for incorporating additional decision-layer criteria in future work, such as predicted toxicity or off-target liability, without altering or retraining the validated biological prediction itself.

Several limitations warrant consideration. Donor-to-donor variability in PBMC functional responses was addressed through fixed donor cohorts per target but was not itself modeled as a source of prediction uncertainty, and future iterations of ClinOracle could incorporate donor-level covariates to better capture this heterogeneity. Validation was limited to five targets within three disease areas; although these were deliberately chosen to span mechanistically distinct immune signaling contexts, broader validation across additional target classes, including non-immune targets, will be necessary to fully establish the generalizability of the platform.

More broadly, these results suggest that the translational attrition long attributed to PK and safety failures may be addressable earlier in the discovery process by explicitly modeling functional activity as a distinct, learnable property rather than assuming it follows automatically from target binding. As patient-derived functional assays become more scalable and standardized, hierarchical frameworks of this kind may offer a general template for AI-guided drug discovery platforms seeking to close the gap between computational prediction and clinical relevance.

## MATERIALS AND METHODS

### Compounds and reagents

All small molecule compounds (**LAG3-A**, **LAG3-B**, **Siglec15-A**, **Siglec15-B**, **STING-A**, **STING-B**, **RIPK1-A**, **RIPK1-B**, **IRAK4-A**, **IRAK4-B**) were obtained from Enamine Ltd. (Kyiv, Ukraine) as part of the Enamine Hit Locator Library (460,160 compounds). Compound identity and purity (>95%) were confirmed prior to biological evaluation. Stock solutions were prepared in DMSO at 10 mM and stored at −80°C. Working solutions were prepared fresh by serial dilution in the appropriate assay buffer immediately before use. Final DMSO concentration in all biological assays did not exceed 0.1% (v/v).

LAG-3 and Siglec-15 ectodomains were expressed in mammalian HEK293 cells to preserve native glycosylation, and for Siglec-15, for sialic-acid-binding function; the STING C-terminal domain was expressed in *E. coli*, and RIPK1 and IRAK4 kinase domains were expressed using a baculovirus/insect cell (Sf9) system. All recombinant proteins were purified by affinity chromatography and used for biophysical binding assays. Anti-LAG-3 antibody (relatlimab, positive control), anti-PD-1 antibody (nivolumab), and isotype control antibodies were purchased from BioXcell and R&D Systems. Cytokine ELISA kits (IFN-γ, IL-2, IFN-α, IL-6, TNF-α, IL-1β, Granzyme B) were purchased from R&D Systems (Minneapolis, MN). All cell culture reagents, including RPMI-1640, FBS, L-glutamine, and penicillin/streptomycin, were purchased from Gibco (Thermo Fisher Scientific, Waltham, MA).

### GNN model architecture and training

#### Dataset preparation

Training datasets for each target (LAG-3, Siglec-15, STING, RIPK1, IRAK4) were assembled from in-house Dianthus MST screening data. Each dataset comprised 25,000 compounds screened from the Training Library with experimentally measured ΔF_norm_ values. Compounds were assigned binary hit labels (1 = hit, 0 = non-hit) using a ΔF_norm_ threshold of ≥20%, corresponding to confirmed target engagement by MST. Molecular graphs were constructed from SMILES strings using RDKit (version 2023.09.1). Node features encoded atomic properties including atomic number, degree, formal charge, number of hydrogens, aromaticity, and hybridization state. Edge features encoded bond properties including bond type, conjugation, ring membership, and stereochemistry. The generation of functional labels for Head 2 training is described below.

#### Model architecture

The ClinOracle GNN employed a shared attentive message-passing neural network (AMPNN) encoder with three graph convolutional layers followed by global attention pooling, feeding two independent classification heads. Attention message passing updated node representations through learned attention coefficients over neighboring node-edge pairs, enabling the model to selectively weight chemically informative interactions. Global attention pooling aggregated node-level representations into a fixed-dimensional molecular embedding h ∈ ℝ^d^ (d = 256), which was passed to each head independently. Each head comprised two fully connected layers (dimensions 256 → 128 → 1) with ReLU activations and dropout (p = 0.2), producing a single output logit.

The binding head predicted P(Binder) from molecular structure across all experimentally screened compounds per target, with a binary binding label (1 if ΔF_norm_ ≥ 20%, 0 otherwise) as the training target. The functional head predicted P(PBMC active | Binder) using a binary functional label (1/0), trained exclusively on compounds with binding label = 1; non-binders were assigned functional mask = 0 and did not contribute to the functional loss. Both heads were trained jointly in a single forward/backward pass per batch: binding loss (binary cross-entropy with logits) was computed over every compound in the batch, and functional loss (binary cross-entropy with logits) was computed only over the confirmed-binder subset of that batch (or omitted for batches containing no confirmed binders). Total loss was computed as binding loss plus functional loss, unweighted (λ = 1). Sigmoid activation was applied to each head’s output logit at inference to yield P(Binder) and P(PBMC active | Binder); training used the raw logits directly with binary cross-entropy with logits loss for numerical stability. The integrated translational probability was computed as P(Translationally active) = P(Binder) × P(PBMC active | Binder). Individual head outputs were not separately calibrated; only the integrated P(Translationally active) score was calibrated via Platt scaling, as described below.

#### Ablation analysis population

For model ablation analyses, a stratified evaluation set comprising 3,500 compounds was assembled from the independent held-out test partitions of all five targets. The evaluation population was modestly enriched for active compounds (5.7% prevalence) to permit stable comparison of alternative model architectures while maintaining a predominantly inactive background. This dataset was completely independent of model training, validation, and calibration. Because PBMC functional measurements were generated only for experimentally confirmed binders, compounds lacking experimental evidence of target binding were considered translationally inactive for endpoint evaluation. Alternative architectures were evaluated on the identical compound set to isolate the contribution of hierarchical modeling to translational prediction performance.

#### Data splitting and validation

For each target, the 25,000-compound dataset was split into training (70%, n = 17,500), validation (10%, n = 2,500), and held-out test sets (20%, n = 5,000) using stratified random splitting to preserve hit rate balance across splits. Model performance was evaluated on the held-out test set using area under the receiver operating characteristic curve (ROC-AUC), with binary labels assigned using the ΔF_norm_ ≥ 20% threshold. Enrichment factors (EF) were calculated at the top 1% of the test set ranked by predicted P(Binder), relative to the baseline hit rate.

#### Prospective virtual screening

Trained GNN models were applied prospectively to the full 460,160-compound Enamine Hit Locator Library, a screening collection entirely independent of the 25,000-compound per-target datasets used for model training, validation, and testing, precluding any compound overlap between the two. SMILES strings for all library compounds were obtained from the Enamine catalog (version March 2026). Molecular graphs were constructed using the same featurization pipeline as training data. Predicted P(Binder) scores were computed for all compounds. The normalized developability score, D, incorporated the quantitative estimate of drug-likeness (QED), Lipinski compliance, Veber compliance, predicted synthetic accessibility (SA), and predicted absorption, metabolism, and excretion (ADME) into a single composite metric. Each compound was assigned this score, calculated as D = 0.30(QED) + 0.15(Lipinski) + 0.15(Veber) + 0.10(SA) + 0.10(Absorption) + 0.10(Metabolism) + 0.10(Excretion), where Absorption, Metabolism, and Excretion were predicted using ADMET-AI (43). Absorption was defined as the predicted probability of high human intestinal absorption; Metabolism was defined as 1 minus the mean predicted inhibition probability across CYP1A2, CYP2C9, CYP2C19, CYP2D6, and CYP3A4, such that higher values indicated lower drug-interaction liability; and Excretion was defined as 1 minus the min-max normalized predicted hepatocyte clearance, such that higher values indicated longer predicted duration of action. Lipinski scores were assigned as 1.0, 0.5, or 0 for zero, one, or at least two violations, respectively; Veber compliance was encoded as 1 or 0; SA scores were rescaled to a 0-1 range (SA = (10 - SA score)/9, clipped to [0,1]) such that higher values indicated greater predicted synthetic accessibility. Each compound’s integrated translational probability, P(Translationally active) = P(Binder) x P(Functional | Binder), was combined with D to yield the ClinOracle Priority Score = P(Translationally active) x D. The top 150 highest-ranked compounds per target were selected for experimental validation, yielding 750 compounds total. Chemical space coverage of ClinOracle-predicted hits was visualized by UMAP dimensionality reduction of Morgan fingerprints (radius = 2, 1,024 bits; RDKit) computed for a representative 50,000-compound subset of the library plus all 750 predicted hits, using Jaccard distance metric.

#### Dianthus microscale thermophoresis (MST)

Binding affinities of GNN-predicted hits were determined using Dianthus MST (NanoTemper Technologies, Munich, Germany). Recombinant target proteins were fluorescently labeled using the RED-NHS labeling kit (NanoTemper Technologies) according to the manufacturer’s instructions. Labeled protein was used at a fixed concentration for each target (10 nM for LAG-3 and STING; 15 nM for Siglec-15; and 20 nM for RIPK1 and IRAK4). Compound stock solutions (10 mM in DMSO) were serially diluted 1:1 in MST buffer (50 mM HEPES pH 7.4, 150 mM NaCl, 0.05% Tween-20, 0.1% BSA) across multiple concentration points ranging from 0.1 nM to 100 μM. Labeled protein and compound dilutions were mixed at a 1:1 ratio and incubated for 30 min at room temperature prior to measurement. MST measurements were performed in premium-coated capillaries using 20% LED power and 40% MST power at 25°C. K_D_ values were determined by sigmoidal dose-response fitting of the fraction bound vs. log[compound] curve. All measurements were performed in quintuplicate (n = 5); K_D_ values are reported as mean ± SD. The two top-ranked ClinOracle candidates per target (A- and B-series) were advanced to orthogonal SPR validation.

#### Surface plasmon resonance (SPR)

SPR binding affinities were measured using a Biacore 8k instrument (from Cytiva) at 25°C. Recombinant target proteins were immobilized on Series S Sensor Chip CM5 (29104988, Cytiva, Marlborough, MA, USA) via standard amine coupling chemistry (EDC/NHS activation). Analyte solutions were prepared by serial dilution in running buffer (10 mM HEPES pH 7.4, 150 mM NaCl, 0.05% Tween-20, 3% DMSO) at multiple concentrations per each tested compound. Samples were injected at a flow rate of 30 μL/min. Surfaces were regenerated between analyte injections using a regeneration buffer. K_D_ values were determined by steady-state affinity analysis using the instrument manufacturer’s software. All measurements were performed in triplicate (n = 3); K_D_ values are reported as mean ± SD. Solvent correction and reference surface subtraction were applied to all sensorgrams.

#### CETSA

Cellular target engagement was assessed by CETSA. Target-specific cell lines, culture conditions, heating temperatures, and ELISA antibodies are summarized in Table S4. Cells were seeded in 6-well plates and cultured in the corresponding medium at 37°C, 5% CO_2_ until 80% confluency. Compound solutions were prepared by serial dilution in culture medium (0.0001-10 μM, final DMSO ≤ 0.1%) and added to cells for 1 hour at 37°C. Following compound treatment, cells were trypsinized, washed twice with PBS, resuspended in PBS, and aliquoted into 0.2 mL PCR tubes. Cell aliquots were heated at a fixed temperature per each protein for 3 minutes using a thermocycler (T100, Bio-Rad), immediately transferred to ice for 3 minutes, and lysed by three freeze-thaw cycles in liquid nitrogen. Lysates were cleared by centrifugation at 20,000 × g for 20 minutes at 4°C. Soluble protein concentrations were quantified as described in Table S4 using antibodies specific to each target protein. Normalized soluble protein (%) was calculated relative to the vehicle-treated control at the same temperature. EC_50_ values were determined by four-parameter sigmoidal (4PL) dose-response fitting using GraphPad Prism (version 10.0). EC_50_ values are reported as mean ± SD (n=5).

#### In vitro ADME profiling

All ADME assays were performed on all 10 ClinOracle-validated compounds (A- and B-series, all five targets) in triplicate (n = 3) unless otherwise stated. Results are presented in Table 1.

#### LogD_7.4_ determination

LogD_7.4_ values were determined by the miniaturized shake-flask method using 1-octanol and PBS (pH 7.4). Compounds were added to a 1:1 mixture of 1-octanol and PBS (pH 7.4) at a final concentration of 10 μM and equilibrated by end-over-end rotation for 2 hours at 25°C. Phases were separated by centrifugation at 3,000 × g for 10 minutes. Compound concentrations in each phase were quantified by LC-MS/MS, and LogD_7.4_ was calculated as log([compound]_octanol_/[compound]_PBS_).

#### Kinetic solubility

Kinetic solubility was determined by the nephelometric method. Compounds were diluted from 10 mM DMSO stocks into PBS (pH 7.4) containing 1% DMSO to achieve a final concentration of 250 μM. After 1 hour incubation at 25°C with shaking, samples were analyzed by UV-vis spectrophotometry at 620 nm to detect precipitation. Solubility was determined by comparison to a calibration curve prepared in 100% DMSO.

#### FaSSIF solubility

Solubility in fasted-state simulated intestinal fluid (FaSSIF, pH 6.8) was determined by the equilibrium shake-flask method. Compounds were added to FaSSIF (prepared according to the BioDis method) at a nominal concentration of 500 μM and equilibrated for 24 hours at 37°C with continuous shaking. Suspensions were filtered through 0.45 μm PVDF membranes, and filtrate concentrations were quantified by LC-MS/MS against calibration standards.

#### PAMPA permeability

Passive membrane permeability was evaluated using the PAMPA Evolution system (pION Inc., Billerica, MA, USA). A phosphatidylcholine-based artificial membrane (1% phosphatidylcholine dissolved in dodecane) was applied to the filter support according to the manufacturer’s instructions. Test compounds were prepared at a final concentration of 10 μM in phosphate-buffered saline (PBS, pH 7.4) containing 0.5% (v/v) DMSO and added to the donor wells, while the acceptor wells contained PBS (pH 7.4). Donor and acceptor plates were assembled and incubated for 4 h at 25°C under static conditions. Following incubation, aliquots from both compartments were analyzed by LC-MS/MS using an internal standard for quantification. The apparent permeability coefficient (P_app_, cm/s) was calculated from the measured donor and acceptor concentrations using the manufacturer’s permeability model, with each compound analyzed in triplicate.

#### Gastrointestinal stability

Stability in simulated gastric fluid (SGF, pH 1.2, pepsin 3.2 mg/mL) and simulated intestinal fluid (SIF, pH 6.8, pancreatin 10 mg/mL) was assessed by incubating compounds at 10 μM for 2 hours at 37°C with continuous shaking. Reactions were quenched with 3 volumes of ice-cold acetonitrile containing internal standard, centrifuged at 3,000 × g for 10 minutes, and supernatants analyzed by LC-MS/MS. Percentage remaining was calculated relative to time-zero controls.

#### Microsomal metabolic stability

Metabolic stability in mouse and human liver microsomes was assessed by incubating compounds at 1 μM with pooled liver microsomes (0.5 mg/mL protein, XenoTech, Kansas City, KS) in 100 mM potassium phosphate buffer (pH 7.4) supplemented with NADPH (1 mM) and MgCl₂ (3 mM) at 37°C. Aliquots were taken at 0, 5, 15, 30, 60, and 100 minutes and reactions terminated by addition of ice-cold acetonitrile (3:1, v/v) containing internal standard. After centrifugation (3,000 × g, 10 min), supernatants were analyzed by LC-MS/MS. Intrinsic clearance (CL_int_) was calculated from the slope of the log-linear decline in compound concentration over time, and t_1/2_ was derived as ln(2)/k_el_. Microsomal half-lives were estimated from the slope of the log-linear concentration-time relationship. For compounds retaining more than 50% of their initial concentration at 100 min, the half-life was not reached and is reported as >100 min.

#### Plasma stability

Plasma stability was assessed by incubating compounds at 1 μM in mouse or human plasma (from Sigma Aldrich) at 37°C with continuous shaking. Aliquots were taken at 0, 5, 15, 30, 60, and 100 minutes and reactions quenched with 3 volumes of ice-cold acetonitrile containing internal standard. PBS stability was assessed identically using PBS (pH 7.4) in place of plasma. Half-lives were calculated as described for microsomal stability. For compounds retaining more than 50% of their initial concentration at 100 min, the half-life was not reached and is reported as >100 min.

#### Plasma protein binding

PPB was determined by rapid equilibrium dialysis (RED) using a 96-well RED device (Thermo Fisher Scientific). Compounds were added to human plasma at 1 μM and dialyzed against PBS (pH 7.4) for 4 hours at 37°C with continuous orbital shaking at 400 rpm. Samples from both compartments were matrix-matched and analyzed by LC-MS/MS. Percentage bound was calculated as [(C_plasma_ − C_PBS_)/C_plasma_] × 100.

#### Cytotoxicity

Cytotoxicity was assessed in WI-38 (normal human lung fibroblasts, ATCC CCL-75) and HS-27 (normal human skin fibroblasts, ATCC CRL-1634) cell lines. Cells were seeded at 5,000 cells/well in 96-well plates and incubated overnight at 37°C, 5% CO_2_. Compounds were added at increasing concentrations ranging (3-fold serial dilution, final DMSO ≤ 0.1%) and incubated for 72 hours. Cell viability was assessed using the CellTiter-Glo luminescent cell viability assay (Promega, Madison, WI). IC_50_ values were determined by four-parameter sigmoidal dose-response fitting using GraphPad Prism (version 10.0).

#### hERG inhibition

hERG potassium channel inhibition was evaluated using an automated whole-cell patch clamp assay (QPatch HTX, Sophion Bioscience) in HEK293 cells stably expressing the human ether-à-go-go-related gene (hERG, Kv11.1) potassium channel. Cells were voltage-clamped according to the manufacturer’s validated protocol, and hERG tail currents were recorded before and after compound application. Test compounds were evaluated in an 8-point concentration-response series (three-fold serial dilutions) prepared in extracellular recording buffer containing a final DMSO concentration of 0.3% (v/v). E-4031 was included as a positive control and vehicle (0.3% DMSO) as a negative control. Peak tail current amplitudes were normalized to vehicle controls, and concentration-response curves were fitted by nonlinear regression using a four-parameter logistic model to determine IC_50_ values. All measurements were performed in triplicate and are reported as mean ± SD.

#### Experimental ADME Score

For inclusion in the integrated translational profile (Figure 5F), a composite Experimental ADME Score was computed for the five A-series lead compounds based on four experimentally determined parameters: human liver microsomal t_1/2_ (metabolic stability), PAMPA P_app_ (passive permeability), kinetic aqueous solubility in 1% DMSO/PBS (developability), and hERG safety margin (hERG IC_50_ / MST K_D_, reflecting the cardiac safety window relative to target affinity). Each parameter was min-max normalized across the five A-series compounds (0 = worst, 1 = best) and the four normalized scores were averaged to yield the composite Experimental ADME Score (range 0-1). This score was not used for compound selection decisions, which were based on individual parameter values; it is presented solely as an integrated summary metric for cross-compound comparison in Figure 5F.

### Patient-derived PBMC functional assays

#### Human PBMC Procurement

Cells were obtained from commercial vendors and were provided to investigators in a de-identified manner. No identifiable donor information was available to investigators. Patient-derived PBMCs were obtained from BioIVT (Westbury, NY) from the following donor cohorts: colorectal cancer patients (n = 10); NSCLC patients (n = 10); SLE patients (n = 10); MS patients (n = 10); and AD patients (n = 10).

#### PBMC isolation and culture

Cryopreserved PBMCs were thawed rapidly at 37°C, washed twice with pre-warmed complete RPMI-1640 (10% FBS, 2 mM L-glutamine, 100 U/mL penicillin, 100 μg/mL streptomycin), and rested for 2 hours at 37°C, 5% CO_2_ prior to use. Cell viability was assessed by trypan blue exclusion and was >90% in all preparations used for assays.

#### Generation of functional labels for Head 2 training

All experimentally confirmed binders identified by MST for each target were subsequently evaluated in the corresponding target-specific patient-derived PBMC functional assay under the same stimulation conditions described below. To enable large-scale training of the functional prediction head, compounds were tested at a single concentration of 1 μM using PBMCs from five independent donors per target, with the primary functional readout specific to each assay. The primary functional readout was IFN-γ for LAG-3 and Siglec-15, IFN-α for STING, TNF-α for RIPK1, and IL-6 for IRAK4. Responses were normalized to vehicle and positive controls. Compounds producing a normalized functional response of ≥30% of the positive-control response without evidence of cytotoxicity (≤20% reduction in viability) were assigned a positive functional label; all others were labeled inactive. These binary labels were used exclusively for training and evaluation of the ClinOracle functional prediction head. The prioritized lead compounds identified by ClinOracle were subsequently evaluated in full dose-response PBMC functional assays as described below.

#### Dose-response functional assay design

For all five PBMC assays presented in Figure 5, compounds were tested across 9 concentration points in the presence of target-specific stimulation, with vehicle control (0.1% DMSO) as the reference condition. Each donor was tested independently with three technical replicates per concentration point (n = 10 donors × 3 replicates = 30 measurements per concentration). Dose-response curves were fitted by four-parameter logistic (4PL) regression using GraphPad Prism (version 10.0). EC_50_ values are reported as mean ± SEM across n = 10 donor-level fitted values. E_max_ values for activation readouts are expressed as fold increase over vehicle; E_max_ values for suppression readouts are expressed as percentage suppression relative to stimulated vehicle.

For suppression readouts (IFN-α, IL-6, TNF-α, NF-κB activation in Panels C, D, and E), data were normalized to the per-donor stimulated vehicle mean prior to averaging across donors, to account for inter-donor variability in baseline inflammatory tone that is independent of compound effect. For activation readouts (IFN-γ, IL-2, tumor lysis, cell viability in Panels A and B), absolute values are presented as the inter-donor spread reflects biologically meaningful differences in T cell responsiveness.

Statistical significance of compound effect vs. vehicle was evaluated at each concentration point using a paired two-tailed t-test (pairing within donor), with Bonferroni correction for multiple comparisons across k = 9 tested concentrations (α_corrected_ = 0.05/9 = 0.00556). Significance thresholds: *p < 0.05, **p < 0.01, ***p< 0.001.

#### LAG-3 functional assay (colorectal cancer PBMCs)

PBMCs from colorectal cancer patients (n = 10) were seeded at 2 × 10^5^ cells/well in 96-well plates pre-coated with anti-CD3 antibody (OKT3, 1 μg/mL) to provide suboptimal T cell stimulation mimicking a tumor-suppressed immune environment. **LAG3-A** was added at concentrations from 1 to 10,000 nM in the presence of soluble anti-CD28 antibody (1 μg/mL) as co-stimulation. After 48 hours at 37°C, 5% CO_2_, supernatants were collected and IFN-γ and IL-2 concentrations were quantified by ELISA (R&D Systems). Anti-LAG-3 antibody (relatlimab, 10 μg/mL) was included as positive control.

#### Siglec-15 functional assay (NSCLC PBMCs)

PBMCs from NSCLC patients (n = 10) were co-cultured with HEK293 cells stably overexpressing human Siglec-15 (AcceGen, Fairfield, NJ, USA) at an effector-to-target ratio of 10:1 in 96-well plates. **Siglec15-A** was added at concentrations from 1 to 10,000 nM. After 24 hours of incubation time, IFN-γ was quantified by ELISA and tumor cell lysis was assessed by LDH release assay (CytoTox 96, Promega), expressed as percentage of maximum lysis achieved by Triton X-100 treatment. Anti-Siglec-15 antibody was included as positive control.

#### STING functional assay (SLE PBMCs)

PBMCs from SLE patients (n = 10) were seeded at 2 × 10^5^ cells/well and stimulated with cGAMP (2’3’-cGAMP, 10 μg/mL, InvivoGen) to activate the STING-IRF3 pathway. **STING-A** was added simultaneously at concentrations from 1 to 10,000 nM. After 24 hours, IFN-α and IL-6 concentrations in supernatants were quantified by ELISA. Data are expressed as percentage of stimulated vehicle per donor.

#### RIPK1 functional assay (MS PBMCs)

PBMCs from MS patients (n = 10) were seeded at 2 × 10^5^ cells/well and treated with TNF-α (20 ng/mL) plus the pan-caspase inhibitor zVAD-FMK (20 μM) to induce necroptosis via the RIPK1-RIPK3-MLKL axis. **RIPK1-A** was added simultaneously at concentrations from 1 to 10,000 nM. After 24 hours, cell viability was assessed by CellTiter-Glo and TNF-α concentration in supernatants was quantified by ELISA. TNF-α suppression data are expressed as percentage of stimulated vehicle per donor; cell viability is expressed as absolute percentage.

#### IRAK4 functional assay (AD PBMCs)

PBMCs from AD patients (n = 10) were seeded at 2 × 10^5^ cells/well and stimulated with LPS (100 ng/mL, Sigma-Aldrich) to activate the TLR4-MyD88-IRAK4 signaling axis. **IRAK4-A** was added simultaneously at concentrations from 1 to 10,000 nM. After 24 hours, IL-6 concentrations in supernatants were quantified by ELISA and NF-κB activation was assessed by luciferase reporter assay. Data are expressed as percentage of stimulated vehicle per donor.

#### In vivo PK studies for LAG3-A

All animal experiments were conducted in accordance with protocols approved by the Institutional Animal Care and Use Committee (IACUC) of Weill Cornell Medicine (Protocol No. 2023-0022). In vivo PK studies were conducted in male C57BL/6 mice (8-10 weeks old; n = 5 per time point). **LAG3-A** was administered by intravenous injection (2 mg/kg) or oral gavage (25 mg/kg) in a formulation consisting of 5% DMSO, 40% PEG400, and 55% sterile saline (v/v/v), prepared fresh prior to dosing. Blood samples were collected at 0.25, 0.5, 1, 2, 4, 8, and 24 h post-dose via tail vein sampling into EDTA-coated tubes. Plasma was isolated by centrifugation at 3,000 × g for 10 min at 4 °C and stored at −80 °C until analysis.

#### In vivo antitumor efficacy (CT26 colorectal tumor model)

All animal experiments were conducted in accordance with protocols approved by the Institutional Animal Care and Use Committee (IACUC) of Weill Cornell Medicine (Protocol No. 2023-0022). Female BALB/c mice (6-8 weeks old; The Jackson Laboratory, USA) were housed under specific pathogen-free conditions with ad libitum access to food and water on a 12-h light/dark cycle and acclimated for at least one week before study initiation.

CT26 murine colorectal carcinoma cells (ATCC CRL-2638) were cultured in RPMI-1640 medium supplemented with 10% fetal bovine serum and 1% penicillin-streptomycin at 37°C in a humidified atmosphere containing 5% CO_2_. Cells were harvested during logarithmic growth (70-80% confluency), washed twice with sterile PBS, and resuspended at 5 × 10^5^ cells/100 μL. Each mouse received a subcutaneous injection of 5 × 10^5^ CT26 cells into the right flank (Day 0).

Tumor growth was monitored three times weekly using digital calipers, and tumor volume was calculated as V = (L × W^2^)/2, where L is the longest tumor diameter and W is the perpendicular diameter. When tumors reached approximately 75-100 mm^3^ (typically Day 7-8), mice were randomized into four treatment groups (n = 11 mice per group) to ensure comparable baseline tumor volumes: (1) vehicle control; (2) **LAG3-A** (25 mg/kg, oral gavage, once daily); (3) anti-mouse PD-1 monoclonal antibody (clone RMP1-14, Bio X Cell; 10 mg/kg, intraperitoneally, twice weekly); and (4) **LAG3-A** (25 mg/kg, oral gavage, once daily) plus anti-mouse PD-1 (10 mg/kg, intraperitoneally, twice weekly). **LAG3-A** was formulated fresh daily in a vehicle consisting of 5% DMSO, 40% PEG400, and 55% sterile saline (v/v/v). The selected dosing regimen was guided by the in vivo PK profile, which maintained plasma concentrations above the biochemical and cellular potency range associated with LAG-3 target engagement. Treatments were administered for 21 consecutive days.

Body weight and clinical condition were monitored throughout the study to assess tolerability. Animals were euthanized upon reaching predefined humane endpoints, including tumor ulceration, impaired mobility, or tumor volume exceeding 2,000 mm^3^, in accordance with institutional guidelines. At study termination, mice were euthanized by CO_2_ inhalation followed by cervical dislocation. Tumors were excised and weighed. Harvested tumors were mechanically dissociated and enzymatically digested with collagenase IV (1 mg/mL, Sigma-Aldrich) and DNase I (100 μg/mL, Sigma-Aldrich) in RPMI-1640 for 30 minutes at 37°C. Single-cell suspensions were filtered through 70 μm cell strainers and washed with PBS. Intratumoral IFN-γ and IL-2 concentrations were quantified using commercial ELISA kits according to the manufacturers’ instructions and expressed as pg/mg total tumor protein.

#### In vivo PK for RIPK1-A

In vivo PK studies were conducted in male C57BL/6 mice (8-10 weeks old; n = 5 per time point). **RIPK1-A** was administered by intravenous injection (2 mg/kg) or oral gavage (25 mg/kg) in a formulation consisting of 5% DMSO, 40% PEG400, and 55% sterile saline (v/v/v), prepared fresh prior to dosing. Blood samples were collected at 0.25, 0.5, 1, 2, 4, 8, and 24 h post-dose via tail vein sampling into EDTA-coated tubes. Plasma was isolated by centrifugation at 3,000 × g for 10 min at 4°C and stored at −80°C until analysis. For CNS exposure analysis, mice were perfused with ice-cold PBS before brain collection to minimize residual blood contamination. Brain tissues were harvested, weighed, and homogenized in ice-cold PBS. Compound concentrations in plasma and brain homogenates were quantified by LC-MS/MS following protein precipitation with acetonitrile containing an internal standard. Chromatographic separation was performed on a C18 column, and analytes were detected in positive electrospray ionization mode using multiple reaction monitoring. Calibration curves were prepared in blank mouse plasma or brain homogenate matrix and used to calculate compound concentrations. PK parameters were calculated by noncompartmental analysis using Phoenix WinNonlin. Brain-to-plasma exposure ratios were calculated from AUC values. Unbound brain-to-plasma ratios (K_p,uu_) were calculated using measured plasma and brain unbound fractions.

#### In vivo efficacy in 5xFAD mice

All procedures were approved by the Institutional Animal Care and Use Committee (IACUC, Protocol #2024-0006). Male and female 5xFAD mice and wild-type littermates (4 months old) were randomized into treatment groups (n = 12 per group; 6 males and 6 females per group).

**RIPK1-A** was administered by oral gavage at 25 mg/kg once daily (QD) for 21 days. Vehicle-treated animals received formulation only. Treatment allocation, behavioral testing, biochemical assays, flow cytometry analysis, and data quantification were performed by investigators blinded to genotype/treatment.

#### Behavioral testing

Behavioral assessments were performed during the light phase following completion of the treatment period by investigators blinded to genotype and treatment. Spatial working memory was first assessed using the Y-maze spontaneous alternation test, followed by evaluation of spatial learning and reference memory using the Morris water maze.

For the Y-maze, mice were placed individually in the center of a three-arm maze and allowed to explore freely for 8 min. Spontaneous alternation was defined as consecutive entries into all three arms without repetition, and the percentage alternation was calculated as the ratio of observed alternations to the maximum possible alternations. Total arm entries were recorded as a measure of locomotor activity.

For the Morris water maze, mice were trained to locate a hidden escape platform submerged beneath the surface of an opaque circular pool using distal visual cues. During the acquisition phase, mice underwent five consecutive days of training, and escape latency to locate the hidden platform was recorded for each trial. Twenty-four hours after the final training session, a probe trial was conducted with the platform removed. Spatial memory retention was assessed by quantifying the percentage of time spent in the target quadrant using an automated video-tracking system.

#### Amyloid quantification

Cortex and hippocampus were microdissected separately, snap-frozen, and homogenized in ice-cold extraction buffer containing protease inhibitors. Tissue homogenates were centrifuged to obtain soluble fractions, and the remaining pellets were further extracted to recover insoluble Aβ. Soluble Aβ was extracted in PBS-based buffer, and insoluble Aβ was extracted from pellets using 5 M guanidine-HCl or 70% formic acid. Aβ42 levels were quantified using a commercially available ELISA kit (Thermo Fisher Scientific; Aβ42, Cat. # KMB3441), according to the manufacturer’s instructions. Concentrations were interpolated from standard curves and normalized to total protein content determined by BCA assay. Data are reported separately for cortex and hippocampus.

#### Cytokine analysis

Brain homogenates were prepared in lysis buffer and cytokine levels (IL-1β, TNF-α) were measured using ELISA kits (R&D Systems, catalog# MLB00C and MTA00B, respectively). Values were normalized to total protein content.

#### Microglial activation

Brain tissues (cortex and hippocampus) were rapidly dissected and processed into single-cell suspensions using the Adult Brain Dissociation Kit (Miltenyi Biotec, Cat. #130-107-677) according to the manufacturer’s protocol. Myelin was removed using Myelin Removal Beads II (Miltenyi Biotec, Cat. #130-096-733). Cells were resuspended in FACS buffer (PBS containing 2% fetal bovine serum and 2 mM EDTA) and incubated with anti-mouse CD16/CD32 Fc block (BioLegend, Cat. #101320) for 10 min at 4 °C to prevent non-specific binding.

Cells were stained for 30 min at 4 °C with fluorophore-conjugated antibodies against CD45 (APC, clone 30-F11, BioLegend, Cat. #103112), CD11b (FITC, clone M1/70, BioLegend, Cat. #101206), TMEM119 (PE, clone 106-6, BioLegend, Cat. #157306), and CD86 (PerCP-Cy5.5, clone GL-1, BioLegend, Cat. #105028). Dead cells were excluded using a viability dye (Zombie NIR, BioLegend, Cat. #423106). Data were acquired on a BD LSRFortessa flow cytometer (BD Biosciences) and analyzed using FlowJo v10. Microglia were defined as live CD45^low^ CD11b^+^ TMEM119^+^ cells. Activated microglia were quantified as the percentage of CD86^+^ cells within the microglial population. A minimum of 50,000 live events per sample were recorded.

#### Statistical analysis

All data are presented as mean ± SD unless otherwise stated. For patient-derived PBMC functional assays, data are presented as mean ± SEM across n = 10 donors, where the donor is the unit of biological replication. Dose-response curves were fitted by four-parameter logistic (4PL) regression; EC_50_ and E_max_ values are reported as mean ± SEM of per-donor fitted values. Statistical significance relative to vehicle was evaluated at each of the nine compound concentrations using paired two-tailed t-tests, with pairing within donor. Bonferroni correction was applied across the nine compound-versus-vehicle comparisons per assay (k = 9; α_corrected_ = 0.05/9 = 0.00556). For between-group comparisons in other contexts, unpaired two-tailed Student’s t-tests were used. For multiple group comparisons, one-way or two-way ANOVA with Tukey’s or Dunnett’s post-hoc tests were applied as appropriate. In vivo tumor growth curves were analyzed by two-way repeated measures ANOVA. Pearson correlation coefficients between GNN-predicted P(Binder) and experimental NFS were computed per target using SciPy. Statistical significance was defined as *p < 0.05, **p < 0.01, ***p < 0.001. All experiments were performed with a minimum of three independent replicates unless otherwise stated.

## Author contributions

The manuscript was written through contributions of all authors. All authors have given approval to the final version of the manuscript. conceptualization: M.T.G. and S.A.A.-R. Methodology: S.A.A.-R. and M.T.G. investigation: S.A.A.-R. and M.T.G. Formal analysis: S.A.A.-R. and M.T.G. visualization: S.A.A.-R. and M.T.G. Supervision: M.T.G. Writing-original draft: M.T.G. and S.A.A.-R. Writing-review and editing: M.T.G. and S.A.A.-R. Data curation: M.T.G. validation: M.T.G. Project administration: M.T.G.

## Data and materials availability

All data needed to evaluate the conclusions in the paper are present inthe paper and/or the Supplementary Materials. The code is available on Zenodo (DOI: https://doi.org/10.5281/zenodo.21736301).

