## Supporting Information for "ClinOracle: Hierarchical AI Prediction of Target Binding and Patient-Derived Functional Activity Across Diverse Therapeutic Targets"

### Contents


Precision-recall curves for the binding head and functional head S3

Correlation between ClinOracle Head 2-predicted functional probability and experimental patient-derived functional efficacy across all five targets S4

Binding-head performance compared against conventional machine learning and representation-learning baselines S5

Calibration of the integrated ClinOracle score before and after Platt scaling S6

MST analysis of **LAG3-A** binding to recombinant murine LAG-3 S7

MST analysis of **RIPK1-A** binding to recombinant murine RIPK1 S7

**RIPK1-A** reduces microglial activation in 5xFAD mice S8

Composition of the screening datasets and train/validation/test partitions for hierarchical model development S9

Quality-control summary for MST and PBMC datasets S10

Computational developability scores for the ten ClinOracle-validated lead compounds S11

Target-specific CETSA experimental conditions S12

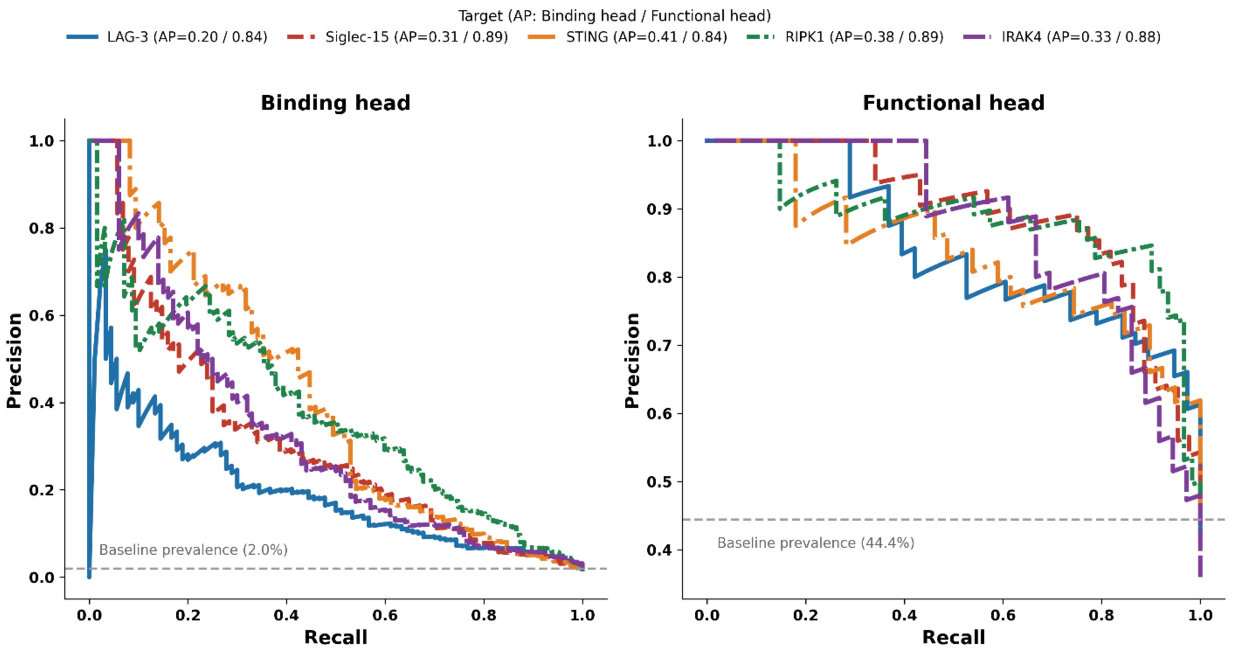


**Figure S1. Precision-recall curves for the binding head and functional head.** Binding-head performance was evaluated across all screened compounds per target (baseline prevalence 2.0%); functional-head performance was evaluated only among experimentally confirmed binders (baseline prevalence 44.4%), reflecting the hierarchical structure of ClinOracle. Average precision (AP) is reported per target for both heads: 0.20-0.41 for the binding head and 0.84-0.89 for the functional head.

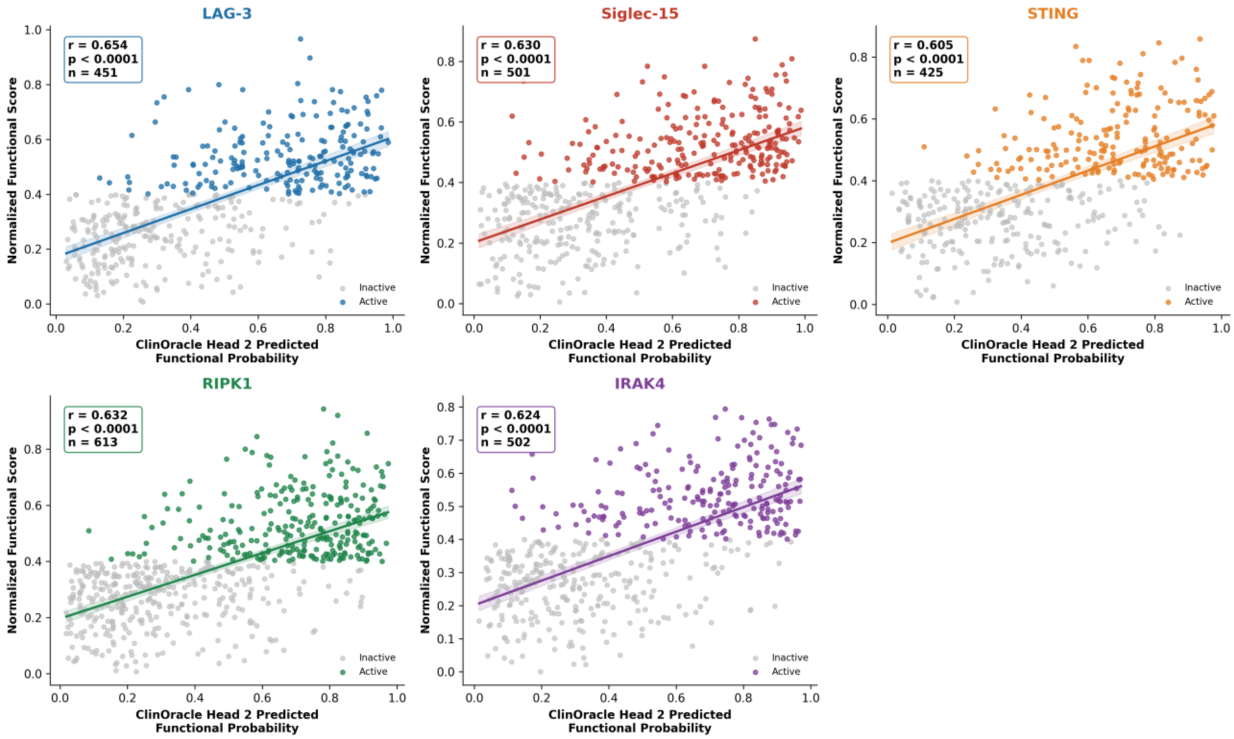


**Figure S2. Correlation between ClinOracle Head 2-predicted functional probability and experimental patient-derived functional efficacy across all five targets.** GNN-predicted functional probabilities from Head 2 (P(PBMC active | Binder)) were significantly correlated with experimentally measured Normalized Functional Scores from patient-derived PBMC functional assays, evaluated across all experimentally confirmed binders per target (n = 425-613 compounds per target). Pearson r ranged from 0.605 (STING) to 0.654 (LAG-3) across all five targets (all $p < 0.0001$). Points are colored by experimentally determined functional status (Active, colored; Inactive, gray). Solid lines show ordinary least-squares regression fits; shaded bands show 95% confidence intervals of the regression estimate.

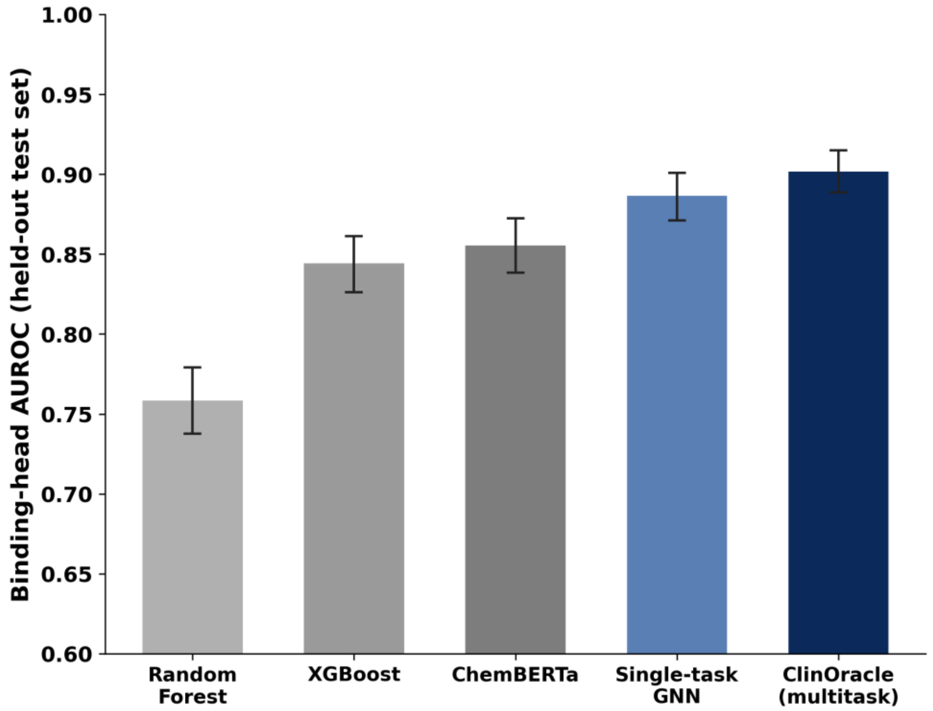


**Figure S3. Binding-head performance compared against conventional machine learning and representation-learning baselines.** Binding-head AUROC on a held-out test set (n = 5,000 compounds per target, pooled across all five targets; n = 25,000 total) for Random Forest (0.758, 95% CI 0.738-0.779), XGBoost (0.844, 0.826-0.861), ChemBERTa (0.856, 0.838-0.872), a single-task GNN (0.887, 0.871-0.901), and the ClinOracle multitask GNN (0.902, 0.889-0.915). Error bars show bootstrapped 95% confidence intervals (2,000 resamples).

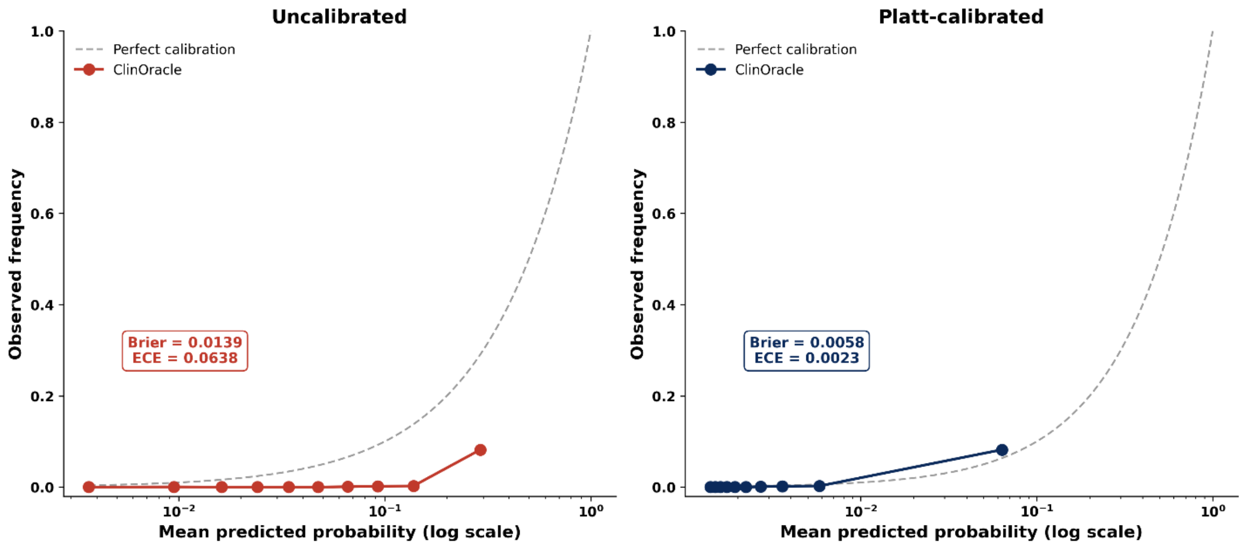


**Figure S4. Calibration of the integrated ClinOracle score before and after Platt scaling.** Mean predicted probabilities are shown on a logarithmic x-axis given the highly skewed score distribution; the dashed curve represents perfect calibration (y = x). A Platt scaling model fitted on the validation split and applied to the held-out test set. Calibration improved substantially (Brier score and expected calibration error shown per panel) while discrimination was fully preserved (AUROC and AUPRC unchanged, as Platt scaling is a monotonic transformation). The uncalibrated score was used for all ranking and compound-prioritization analyses in the main text; the calibrated score provides an interpretable probability estimate where required.

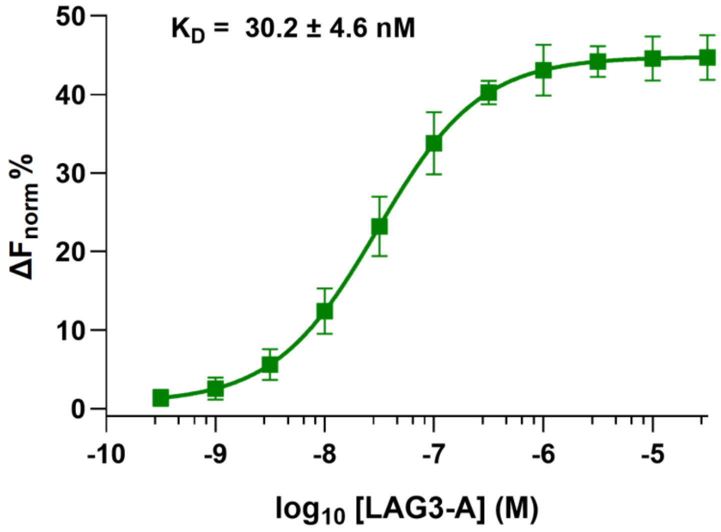


**Figure S5. MST analysis of LAG3-A binding to recombinant murine LAG-3.** Direct interaction was assessed under solution-phase conditions, revealing concentration-dependent binding of **LAG3-A** to murine LAG-3. Data are presented as mean ± SD (n = 5).

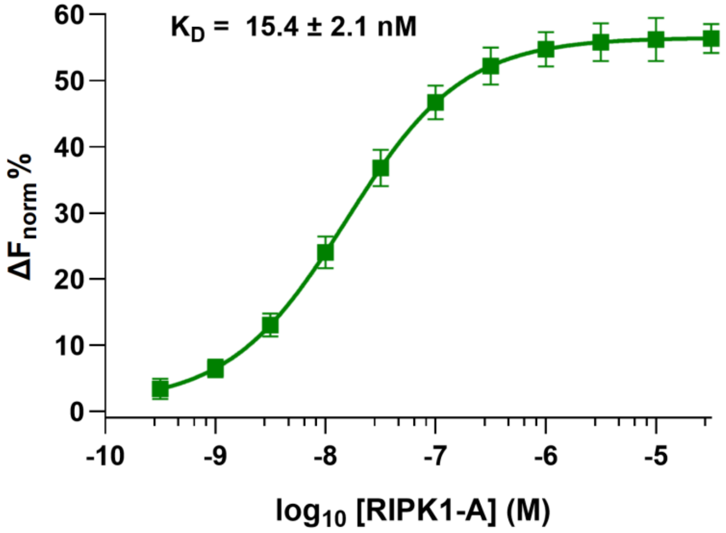


**Figure S6. MST analysis of RIPK1-A binding to recombinant murine RIPK1.** Direct interaction was assessed under solution-phase conditions, revealing concentration-dependent binding of **RIPK1-A** to murine RIPK1. Data are presented as mean ± SD (n = 5).

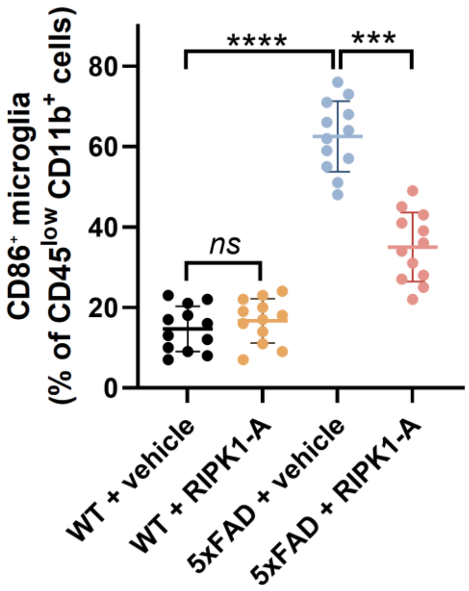


**Figure S7. RIPK1-A reduces microglial activation in 5xFAD mice.** Microglial activation was quantified as the percentage of $CD86^{+}$ cells within the $CD45^{low}CD11b^{+}$ microglial population. Vehicle-treated 5xFAD mice exhibited a marked increase in $CD86^{+}$ microglia relative to WT controls, which was significantly attenuated by **RIPK1-A** treatment. **RIPK1-A** did not significantly alter microglial activation in WT mice. Data are presented as mean ± SD (n = 12 mice per group). Comparisons were performed using one-way ANOVA followed by Tukey's multiple-comparisons test. ***$p < 0.001$, ****$p < 0.0001$; ns, not significant.

**Table S1. Composition of the screening datasets and train/validation/test partitions for hierarchical model development.** Each target comprised 25,000 screened compounds. Each target-level dataset was partitioned once at the compound level into training, validation, and held-out test sets (70%/10%/20%). The same partition assignments were used for both prediction heads during joint model training and evaluation. Binding loss was computed across all compounds within each partition, whereas functional loss was computed only for experimentally confirmed binders within the corresponding partition. Thus, confirmed binders evaluated by the functional head inherited their partition assignments from the complete target-level binding dataset, and no validation or test compound contributed to model training through either prediction head. The functional train, validation, and test counts represent the numbers of confirmed binders falling within these inherited compound-level partitions. No compound appeared in more than one partition within a given prediction task.

| Target | Total screened | Confirmed binders (hit rate) | Functional split (Train/Val/Test) | Functionally active (rate) |
|---|---|---|---|---|
| **LAG-3** | 25,000 | 451 (1.80%) | 314 / 47 / 90 | 203 (45.0%) |
| **Siglec-15** | 25,000 | 501 (2.00%) | 367 / 46 / 88 | 243 (48.5%) |
| **STING** | 25,000 | 425 (1.70%) | 311 / 29 / 85 | 190 (44.7%) |
| **RIPK1** | 25,000 | 613 (2.45%) | 424 / 62 / 127 | 265 (43.2%) |
| **IRAK4** | 25,000 | 502 (2.01%) | 350 / 52 / 100 | 212 (42.2%) |

**Table S2. Quality-control summary for MST and PBMC datasets.**
The number of compounds passing automated quality control (Pass) and requiring manual review (Review) is shown for each target. All analyses reported in the manuscript were performed using the complete dataset, including both Pass- and Review-flagged compounds. As a representative sensitivity analysis, exclusion of Review-flagged compounds for the STING dataset produced a negligible change in binding-head performance (AUROC, 0.9106 vs. 0.9105), indicating that model performance is robust to quality-control filtering. Given the very low proportion of Review-flagged compounds across all targets (0.12-0.17% for MST; 0.20-0.89% for PBMC), additional target-specific sensitivity analyses were not performed.

| Target | MST Pass | MST Review (%) | PBMC Pass | PBMC Review (%) |
|---|---|---|---|---|
| **LAG-3** | 24,970 | 30 (0.12%) | 447 | 4 (0.89%) |
| **Siglec-15** | 24,959 | 41 (0.16%) | 499 | 2 (0.40%) |
| **STING** | 24,961 | 39 (0.16%) | 423 | 2 (0.47%) |
| **RIPK1** | 24,957 | 43 (0.17%) | 610 | 3 (0.49%) |
| **IRAK4** | 24,969 | 31 (0.12%) | 501 | 1 (0.20%) |

**Table S3. Computational developability scores for the ten ClinOracle-validated lead compounds.**

| Compound | MW | LogP | HBD | HBA | TPSA (Å²) | Rot. Bonds | QED | Lipinski Violations | Veber | M | E | D |
|---|---|---|---|---|---|---|---|---|---|---|---|---|
| **LAG3-A** | 394.5 | 2.27 | 2 | 5 | 95.0 | 5 | 0.692 | 0 | Pass | 0.784 | 0.729 | 0.836 |
| **LAG3-B** | 322.4 | 3.22 | 1 | 3 | 61.9 | 3 | 0.944 | 0 | Pass | 0.504 | 0.601 | 0.870 |
| **Siglec15-A** | 341.5 | 2.45 | 1 | 4 | 65.1 | 5 | 0.907 | 0 | Pass | 0.986 | 1.000 | 0.955 |
| **Siglec15-B** | 279.3 | 1.58 | 2 | 5 | 84.1 | 5 | 0.814 | 0 | Pass | 0.635 | 0.000 | 0.792 |
| **STING-A** | 373.4 | 3.02 | 2 | 4 | 67.8 | 6 | 0.817 | 0 | Pass | 0.624 | 0.572 | 0.847 |
| **STING-B** | 325.4 | 3.91 | 1 | 3 | 43.0 | 5 | 0.773 | 0 | Pass | 0.350 | 0.528 | 0.803 |
| **RIPK1-A** | 379.5 | 2.62 | 2 | 4 | 60.5 | 5 | 0.838 | 0 | Pass | 0.761 | 0.872 | 0.898 |
| **RIPK1-B** | 366.5 | 4.83 | 1 | 4 | 51.2 | 6 | 0.680 | 0 | Pass | 0.323 | 0.083 | 0.733 |
| **IRAK4-A** | 403.5 | 2.91 | 1 | 5 | 72.3 | 6 | 0.685 | 0 | Pass | 0.419 | 0.351 | 0.768 |
| **IRAK4-B** | 342.4 | 2.30 | 2 | 6 | 92.3 | 4 | 0.886 | 0 | Pass | 0.909 | 0.700 | 0.910 |

MW, molecular weight (g/mol); LogP, calculated octanol-water partition coefficient (Crippen); HBD/HBA, hydrogen bond donors/acceptors; TPSA, topological polar surface area; QED, quantitative estimate of drug-likeness; Lipinski Violations, count of Rule-of-Five violations; Veber, pass/fail on rotatable bonds ≤10 and TPSA ≤140; M, 1 − mean predicted CYP1A2/2C9/2C19/2D6/3A4 inhibition probability; E, 1 − min-max normalized predicted hepatocyte clearance; D, normalized developability score (D = 0.30(QED) + 0.15(Lipinski) + 0.15(Veber) + 0.10(SA) + 0.10(A) + 0.10(M) + 0.10(E), where SA is a synthetic-accessibility term and A is predicted intestinal absorption probability). The Absorption term is omitted from this table as all 10 compounds scored uniformly high (A > 0.999), reflecting their favorable TPSA relative to the absorption-limiting threshold; A retains discriminating value across the broader screened library. QED/Lipinski/Veber/SA computed using RDKit (version 2023.09.1); A/M/E predicted using ADMET-AI.

**Table S4. Target-specific CETSA experimental conditions.**

| Target | Cell line | Culture medium | CETSA temperature (°C) | Protein quantification |
|---|---|---|---|---|
| **LAG-3** | Raji-hLAG-3 cells (from InvivoGen) | RPMI-1640 + 10% FBS + 1% Penicillin/Streptomycin | 50 | Human LAG-3 ELISA<br>(R&D Systems, Catalog # DY2319B) |
| **Siglec-15** | HEK293 cells stably overexpressing human Siglec-15 (from AcceGen) | DMEM (high glucose) + 10% FBS + 1% Penicillin/Streptomycin | 52 | Human Siglec-15 ELISA<br>(AssayGenie, Catalog# AEKE00006) |
| **STING** | HEK293 cells stably overexpressing human STING (from AcceGen) | DMEM (high glucose) + 10% FBS + 1% Penicillin/Streptomycin | 50 | Human STING ELISA<br>(abcam, Catalog#ab315058) |
| **RIPK1** | HEK293 cells expressing human RIPK1 | DMEM (high glucose) + 10% FBS + 1% Penicillin/Streptomycin | 49 | Human RIPK1 ELISA<br>(Novus Biologicals, Catalog#NBP3-18223) |
| **IRAK4** | HEK293 cells expressing human IRAK4 | DMEM (high glucose) + 10% FBS + 1% Penicillin/Streptomycin | 51 | Human IRAK4 ELISA<br>(abcam, Catalog#ab213472) |
